# Hydrogen exchange of chemoreceptors in functional complexes suggests protein stabilization mediates long-range allosteric coupling

**DOI:** 10.1101/676783

**Authors:** Xuni Li, Stephen J. Eyles, Lynmarie K. Thompson

## Abstract

Bacterial chemotaxis receptors form extended hexagonal arrays that integrate and amplify signals to control swimming behavior. Transmembrane signaling begins with a 2 Å ligand-induced displacement of an alpha helix in the periplasmic and transmembrane domains, but it is not known how the cytoplasmic domain propagates the signal an additional 200 Å to control the kinase CheA bound to the membrane-distal tip of the receptor. The receptor cytoplasmic domain has previously been shown to be highly dynamic as both a cytoplasmic fragment (CF) and within the intact chemoreceptor; modulation of its dynamics are thought to play a key role in signal propagation. Hydrogen deuterium exchange mass spectrometry (HDX-MS) of functional complexes of CF, CheA, and CheW bound to vesicles in native-like arrays reveals that the CF is well-ordered only in its protein interaction region where it binds CheA and CheW. Rapid exchange is observed throughout the rest of the CF, with both uncorrelated (EX2) and correlated (EX1) exchange patterns, suggesting the receptor cytoplasmic domain retains disorder even within functional complexes. HDX rates are increased by inputs that favor the kinase-off state. We propose that chemoreceptors achieve long-range allosteric control of the kinase through a coupled equilibrium: CheA binding in a kinase-on conformation stabilizes the cytoplasmic domain, and signaling inputs that destabilize this domain (ligand binding and demethylation) disfavor CheA binding such that it loses key contacts and reverts to a kinase-off state. This study reveals the mechanistic role of an intrinsically disordered region of a transmembrane receptor in long-range allostery.

## Introduction

Bacterial chemotaxis is employed by both beneficial and pathogenic bacteria to direct their swimming towards favorable environments. Chemotaxis receptors provide the sensory input for this process by binding ligands in their periplasmic domain and transmitting a signal that controls the activity of a kinase (CheA) bound to the receptor cytoplasmic domain. CheA transfers phosphoryl groups to response regulator proteins that control the rotation of the flagellar motor and also control the methylation state of the receptor (adaptation). The kinase activity of CheA is controlled by both the ligand occupancy of the receptor and its adaptation state (adaptation involves methylation and demethylation of 4-5 glutamic acid residues within the cytoplasmic domain). When an attractant ligand binds to the receptor, the tumbling frequency of the cell initially decreases, and then adaptation returns it to its previous level; this enables the bacterium to bias its random walk towards favorable environments. The network of protein interactions involved in chemotaxis is well-understood, but the mechanism of receptor control of the kinase remains elusive. Understanding this mechanism could provide a foundation for harnessing chemotaxis for sensing or delivery applications, or for inhibiting chemotaxis with novel antibiotics.

Extensive studies of chemotaxis receptors have yielded structural models and insights into the signaling mechanism. Receptor complexes with CheA and a coupling protein CheW form remarkable hexagonal arrays that extend ∼200 nm in the membrane of the cell. These arrays consist of trimers of receptor dimers that are bridged at their membrane-distal cytoplasmic tips by a “baseplate” consisting of alternating CheA/CheW rings and CheW-only rings (1, 2). The chemotaxis receptors are also quite long (∼300 Å for the well-studied *E. coli* receptors), and thus ligand binding in the periplasm must induce a long-range allosteric change to control the kinase. It is widely accepted that signaling begins with a ligand-induced 2 Å piston displacement of a receptor helix that extends through the periplasmic and transmembrane domains (3). A key question in the field is how the signal then propagates an additional ∼200 Å through the cytoplasmic domain to control kinase activity.

Previous studies have shown that the receptor cytoplasmic domain is highly dynamic, at least in the absence of its cytoplasmic binding partners CheA and CheW. A cytoplasmic fragment (CF)^1^ of the Asp receptor, containing all but the membrane-proximal HAMP domain, has been shown to be highly dynamic, having an alpha helical secondary structure but a fluctuating tertiary structure (4). Furthermore, the cytoplasmic domain of the intact receptor retains these dynamics, based on its promiscuous disulfide crosslinking patterns (5). Finally, inverse modulation of dynamics of different regions has been suggested to play an important role in signal propagation through this domain. In particular, the ligand-bound, kinase-off state of the receptor is proposed to have a stabilized HAMP domain, destabilized methylation region, and stabilized protein interaction region relative to the kinase-on state (6, 7).

We have used hydrogen-deuterium exchange mass spectrometry (HDX-MS) to investigate changes in structure and dynamics throughout most of the cytoplasmic domain of the receptor. This requires preparation of homogeneous, functional complexes of the receptor with CheA and CheW. Such homogeneity has not yet been achieved for the intact receptor, due to the challenge of controlling the orientation of insertion of the intact receptor into membrane vesicles or nanodiscs. Thus, we chose to assemble the aspartate receptor CF into native-like, functional complexes with CheA and CheW, using a vesicle templating method developed by Weis and coworkers (8). CF in solution is not functional: it does not activate CheA and is not efficiently methylated. Binding of the His-tagged CF to the surface of large unilamellar vesicles containing a mixture of DOPC and DGS-NTA(Ni), in the presence of CheA and CheW, restores function (CF becomes a good substrate for methylation and activates CheA) (8) and also yields native-like hexagonal arrays (9). We have previously used HDX-MS on such complexes to show that incorporation of CF into functional complexes reduces its rapid hydrogen exchange (10), and to measure differences in the exchange properties of functionally important receptor subdomains (11).

We have now conducted an extensive analysis of HDX of CF in functional complexes, with improved samples (more homogeneous CF), better MS data (about three times as many peptides for each of three states), and more thorough data analysis of HDX (of overall uptake, EX1, and EX2 exchange). This study is the first to document widespread EX1 throughout fully complexed CF and to delineate the boundaries of regions with rapid vs very slow HDX, corresponding to previously identified regions of CF. The HDX properties demonstrate that CF within functional arrays is quite stable in its small protein interaction region (where the arrays show greatest order) and marginally stable elsewhere. We propose that signaling inputs (ligand binding and demethylation) decrease the stability of the CF, which in turn disrupts a key contact with CheA to turn off its kinase activity. This novel mechanism employs protein stabilization to achieve integration of multiple inputs for long-range allosteric control of the kinase activity.

## Results

### HDX-MS approach for investigating properties of receptor cytoplasmic domain within functional complexes

For HDX-MS experiments, native-like complexes of CF, CheA, and CheW are assembled on vesicles and then tested for kinase activity and incorporation of proteins into the array. Homogeneity is maximized by (1) using excess CheA and CheW to drive all of the CF into functional complexes, and (2) using a high enough lipid concentration to accommodate all of the CF in functional complexes, based on the known geometry of the hexagonal array observed in cells. Measurements of kinase activity and protein binding (sedimentation assay) are performed before hydrogen exchange and after exchange for 16 hours. These measurements demonstrate that complexes have high kinase activity, protein stoichiometries comparable to native arrays, and no significant loss of activity or proteins throughout the HDX time course (Figure 1 and Table S1). After 16 hours, both high activity (100% for CF4Q and 63% for CF4E) and native-like protein stoichiometries are retained. Thus, the properties observed by HDX-MS are those of the cytoplasmic domain of the receptor within functional signaling complexes.

**Figure 1.**
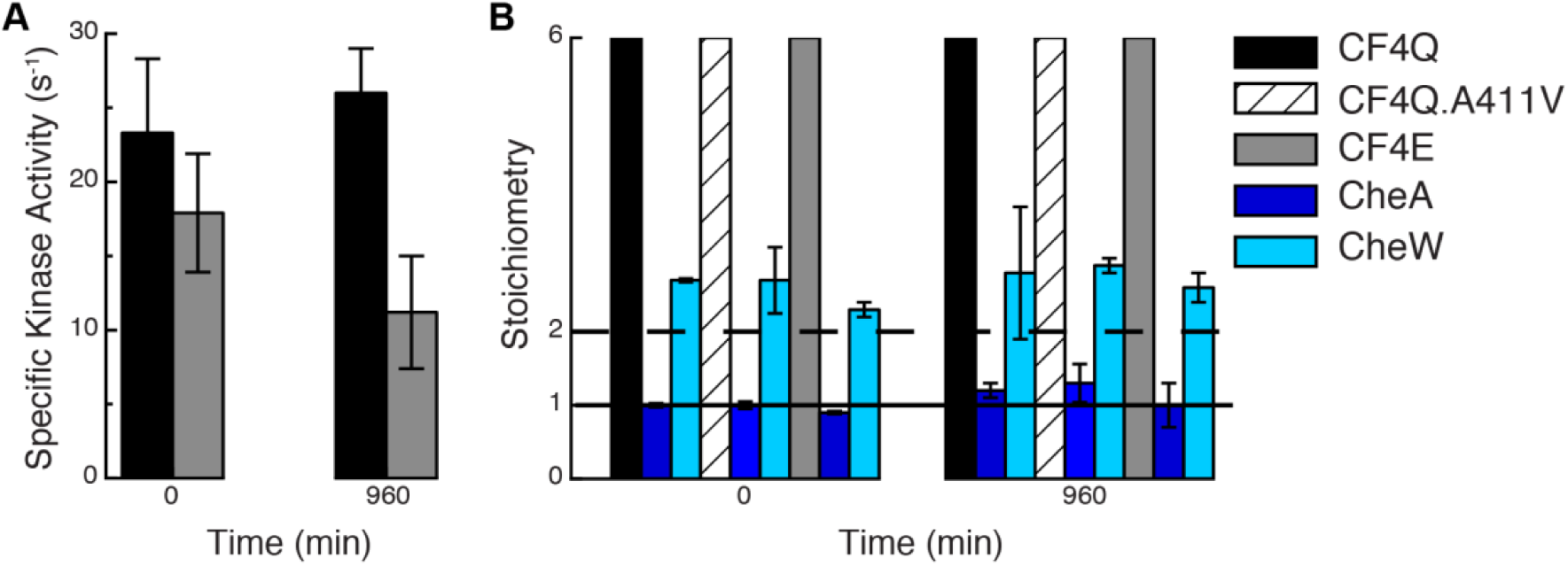
CF complexes retain native kinase activity and protein stoichiometry over the time-course of deuterium labeling. (A) Specific activity of CheA kinase assembled into native-like complexes with CheW and His-tagged CF4Q (black) or CF4E (gray) bound to vesicles. For each complex, kinase activity is high and does not change significantly between 0 and 960 min (=16 hr). Activities are averages of four replicates measured on two different days; error bars indicate ± one standard deviation. (B) Protein stoichiometries, determined as the ratio of proteins in the sedimented complexes (SDS-PAGE band intensities), within complexes of CF4Q and CF4E (both kinase-on) and CF4Q.A411V (kinase-off). Sedimented CF (24-29 µM) was set to 6 for calculations of the ratio of CF to CheA (blue) and CheW (cyan). Horizontal lines correspond to the levels of CheA (solid line) and CheW (dashed line) observed in native arrays.

To investigate the differences in HDX properties between signaling states we examined the effect of methylation and of a mutation that turns off the kinase. CF4E, with Glu at all four methylation sites, represents the unmethylated state. CF4Q represents the methylated state, because amidation at the methylation sites has been shown to have the same effect on signaling as methylation (12). To represent the kinase-off state we used CF4Q.A411V. The A411V mutation in the intact receptor does not interfere with assembly of complexes with CheA and CheW, but locks the receptor into a kinase-off state (13). We have shown that CF4E.A411V exhibits enhanced methylation rates, as expected for the kinase-off state (14). This inverse effect on activity (decreased kinase and increased methylation activity) is a key indicator that a mutation shifts the receptor between its physiologically relevant signaling states, rather than disrupting its activity in some other fashion.

Current sample conditions represent a significant improvement over our previous HDX study. The samples used previously were similar, but CF4E complexes were assembled at low density (high vesicle surface area, kinase-off state) and high density (low vesicle surface area, kinase-on state (15)). We have since realized that the high density conditions did not have sufficient vesicle surface area to allow all of the CF to assemble into functional complexes with CheA and CheW. We estimate that the limited vesicle surface area of the high density samples led to 47% of the CF4E being bound within close-packed CF-only regions rather than forming the hexagonal array with CheA and CheW. For the current study all samples are assembled at an intermediate surface density so that both CF4Q and CF4E are kinase-on, and the surface area is high enough to accommodate >90% of bound CF in functional complexes. We have also used a mutant to control signaling state so that we can assemble all complexes at the same density. This avoids possible perturbations to HDX via changes in solvent exposure. Finally, we have also used two-fold higher concentrations of CheA and CheW to drive all of the CF into functional complexes, as demonstrated by the 6 CF: 1 CheA: 2 CheW stoichiometry (Figure 1).

The HDX experiment is initiated by a rapid transfer of the complex from H_2_O to D_2_O. This is typically achieved in HDX studies by a 15-20-fold dilution into D_2_O, but such dilution of a protein complex could cause dissociation during the exchange time course. Therefore, exchange is initiated without significant dilution using a spin column, pre-equilibrated with deuterated kinase buffer at 25 °C. For each exchange time point, an aliquot is removed, added to pre-chilled quench buffer, and immediately flash-frozen. For MS analysis, each sample of the hydrogen exchange series is thawed and digested with pepsin at 0°C, followed by LC-ESI-MS analysis on a Waters Synapt G2Si Q-TOF mass spectrometer. At least one undeuterated sample and one fully deuterated sample (prepared by reversible heat denaturation of CF in deuterated buffer) are included as controls on each day of mass spectrometry; the fully deuterated sample (which undergoes back exchange equivalent to that of all of the samples) provides a measure of exchange endpoint.

The peptides shown in Figure S1 were identified in the mass spectra of the undeuterated (0 min time point) samples, acquired in collision-induced dissociation mode for peptide sequencing. We identified a total of 86 CF peptides, 183 CheA peptides, and 52 CheW peptides (on average for the different samples), providing an average CF peptide coverage of 80-88% and amino acid redundancy of 2.6-3.4 for the functional complexes of CF4Q, CF4Q.A411V and CF4E. The peptides of the completely exchanged CF4Q control sample exhibited 11-43% back exchange for our protocol (average 25%), with little variation (1%) for experiments performed on different days. Compared to our prior HDX study (11), the use of improved instrumentation including ion mobility separation on the Waters Synapt G2Si led to better reproducibility and resolution, as well as software (DynamX 3.0) enabling us to identify more and shorter peptides that better localize the HDX properties in the CF.

### HDX rates indicate CF has very stable protein interfaces but is otherwise highly dynamic

Figure 2 illustrates hydrogen exchange throughout CF4Q within functional, kinase-on complexes assembled with CheA and CheW on vesicles. The mass spectra for many peptides exhibit bimodal isotopic distributions (discussed below), but some key features of the exchange properties can be deduced from analysis of the centroid mass (intensity-weighted average mass). The percent deuterium uptake of each peptic peptide is calculated by subtracting the centroid mass of the undeuterated sample from that of the deuterated sample at each time point, and then dividing by the uptake of the fully deuterated control sample (complete exchange). Deuterium uptake data are shown in Figure 2A for representative peptides, and illustrate the 80% sequence coverage (all but the black regions). Uptake data for the full set of all peptides from functional complexes of CF4Q, CF4Q.A411V, and CF4E, as well as CF4Q alone, are shown in Figure S2. Peptide numbering throughout the paper corresponds to the sequence of the intact *E. coli* Asp receptor.

**Figure 2.**
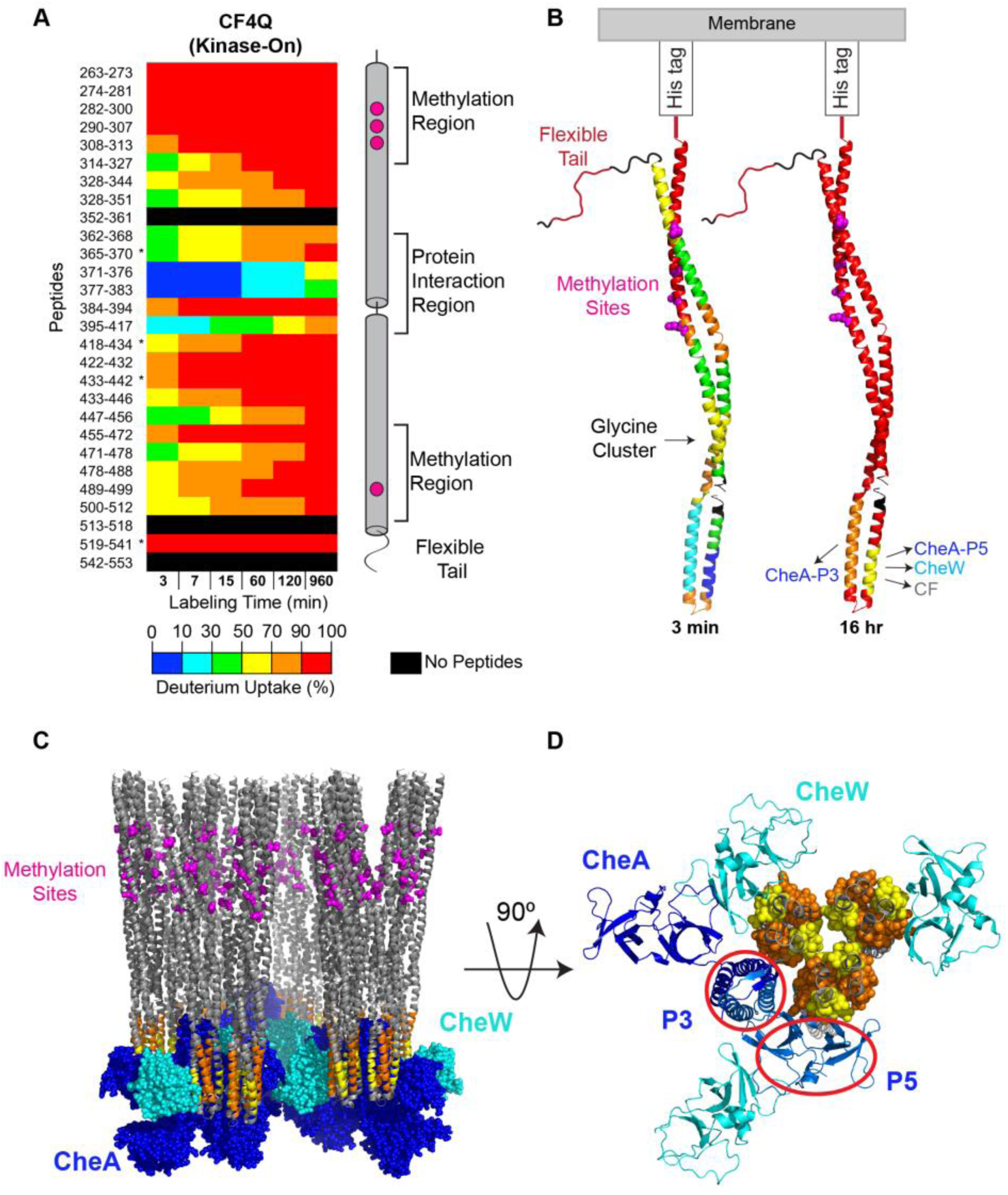
Overview of hydrogen exchange properties of CF4Q in functional complexes assembled with CheA and CheW on vesicles. (A) Deuterium uptake vs time for representative CF peptides covering the entire sequence, except segments shown in black. Percent deuterium uptake of peptides at each time point is represented in rainbow colors from blue to red, indicating low to high deuterium incorporation. Data are averages of 2 replicates, except for *peptides that were missing in one data set. Secondary structure elements of CF are shown on the right, with methylation sites (magenta) and different regions indicated. (B) CF structure is colored according to the percent deuterium uptake at the first and last time points, providing an overview of initial exchange rates (3 min) and of incomplete exchange at long times (960 min = 16 hr). For clarity, only a monomer is shown (PDB file 1qu7 in all figures). Colors are chosen based on averaging the data for overlapping peptides (see black boxes in Figure S2). Segments with incomplete exchange at 16 hours correspond to sites of interactions with the indicated proteins. (C) Side and (D) bottom-up views of a structural model of functional complexes of CF, CheA, and CheW. CF is colored gray, except peptides showing incomplete exchange at 16 hr are colored as in (B). The orange peptide (395-417) is at the interface with the P3 dimerization domain of CheA (red circle). The yellow peptides (371-376, 377-383) are at the interface with the P5 domain of CheA (red oval), or with CheW (cyan), or with another CF dimer in the center of the receptor trimer-of-dimers. Model of complex shown in C and D is based on the array model of Briegel, Crane, and Jensen (1) with an additional CheW positioned via superposition of the neighboring CF dimer tip with array model 3AJ6 (31).

The initial deuterium uptake for the first 3 min time point (Fig 2A & B) varies widely from >90% (red) near the N and C termini to <10% (blue) near the membrane-distal tip. Peptides near the N and C termini exhibit very fast exchange, with 90-100% deuterium uptake in the first 3 minutes, suggesting these regions are highly flexible and solvent exposed. These findings are consistent with previous hydrogen exchange mass spectrometry (11) and NMR studies (14) on similar samples of functional complexes, and with electron paramagnetic resonance (EPR) on spin-labelled intact receptors in nanodiscs (16). The C-terminal tail is thought to be an unstructured, flexible tether for binding the methylation and demethylation enzymes (17). Its very fast hydrogen exchange is comparable to that of residues 263-307, which follow the membrane-anchored N-terminal His tag (residues 248-253). This suggests that MH1 (methylation helix 1, 263-318) is also an unstructured, flexible segment.

At the other extreme, the slowest hydrogen exchange occurs near the membrane-distal cytoplasmic tip of the CF, in the protein interaction region. With the exception of the 384-394 peptide at the hairpin tip of the CF, which exhibits fast exchange (80% at 3 min), residues 369–417 exhibit only 2–17% exchange in the first 3 minutes. The slow exchange in this region on both sides of hairpin tip continues throughout the time course, resulting in incomplete exchange at the final time point (16 hours) for these peptides. This slow exchange is likely due to protein-protein interactions in this region, as illustrated in a structural model of the CF array (1) in Figures 2C and 2D, with the regions of incomplete exchange colored by their uptake at 16 hours (as in Figure 2B) and the rest of the CF colored gray. The 369-383 region (yellow, see Figure S2) is involved in protein interactions with the P5 domain of CheA (red oval), with CheW (homologous to P5), and with CF at the center of the trimer-of-dimers. Similarly, the 395-417 region (orange) is involved in protein interactions with the P3 dimerization domain of CheA (red circle).

Thus the 310 amino acid CF exhibits very slow exchange for 12% (38) of its residues due to stable protein interactions, and very fast exchange for 23% (71) of its residues that are likely unstructured or rapidly sampling multiple structures. The remaining 154 residues observed by HDX (50% of the CF) exhibit fast exchange, with ≥50% uptake in 15 min. These overall properties observed for the kinase-on arrays of CF4Q, very fast exchange for peptides near the N and C termini, slow exchange for peptides in the protein interaction region (with the exception of the hairpin tip itself), and fast exchange everywhere else, are also observed in functional arrays of CF4Q.A411V (kinase-off mutant) and CF4E (other methylation extreme), as shown in Figure S2. For comparison, the hydrogen exchange time course for CF4Q alone (in the absence of vesicles, CheA and CheW; right column in Figure S2) shows essentially complete exchange throughout the protein in 3 minutes. Thus, although formation of functional complexes stabilizes CF, the majority (73%) of CF exhibits fast to very fast HDX, suggesting a dynamic/open structure within all three types of functional complexes. Interestingly, mass spectra of all of the functional complexes also exhibit widespread bimodal isotope distributions, as discussed below.

### Widespread correlated exchange suggests that CF populates an open/dynamic structure in slow exchange with a more folded species

HDX data for many CF peptides exhibit bimodal isotopic distributions, with one isotopic distribution at low m/z that moves gradually to higher mass with time (uncorrelated exchange), and another isotopic distribution at the final exchange m/z position that increases in intensity with time (correlated exchange). Uncorrelated exchange is often observed in proteins under native conditions, where the refolding rate of the protein is faster than the exchange (known as the EX2 limit), so the protein must undergo multiple unfolding/refolding events before HDX is complete. Correlated exchange is observed when the protein refolding rate is slower than the exchange (known as the EX1 limit): the protein populates an unfolded or open state which undergoes complete exchange before refolding. Although the HDX patterns observed for CF include bimodal patterns indicative of correlated exchange (EX1), the deuterium uptake analysis of the HDX data described above was performed with the instrument software, which calculates the centroid of each spectrum (intensity-weighted mass average) and thus ignores the bimodal patterns and the underlying EX1 process. We pursued additional analysis of these bimodal patterns to gain insight into the unfolded or open state and how it relates to methylation and signaling states.

Peptides with HDX mass spectra with bimodal distributions are observed throughout the CF in all functional complexes with CheA and CheW. However, the HDX spectra do not follow an idealized EX1 pattern, with mass envelopes only at zero exchange and complete exchange. Instead a simultaneous EX2 exchange occurs, as expected for “realistic” EX1 (18): the lower m/z envelope shifts gradually to higher m/z, merging with the envelope at the fully exchanged position. Although some HDX series of mass spectra can be deconvoluted into the EX1 and EX2 uptake profiles, this is not possible in many cases due to the incomplete resolution of the bimodal distributions. We implemented an efficient method to estimate t_1/2_ of the EX2 and EX1 exchange processes for each peptide that exhibited HDX with bimodal isotopic distributions. For the EX2 exchange, the t_1/2_ was estimated as the time at which the tallest peak of the gradually shifting isotopic distribution reached an m/z value halfway between the unexchanged and fully exchanged control spectra. Since EX1 exchange populates the fully exchanged isotopic distribution (at high m/z), this process reaches 50% completion when this high m/z mass envelope contains 50% of the intensity in the spectrum. Thus, at t_1/2_ for the EX1 process the bimodal distribution should be symmetric, with equal intensities in the mass envelopes at the low m/z position and at the fully exchanged position. The symmetry of the pattern is an especially useful indicator when the mass envelopes have merged, at which point it is difficult to estimate the intensity in the high m/z mass envelope.

To validate our method, we compare the t_1/2_ values estimated visually with the results of deconvolution analysis with HX-Express (19). Figure S3 shows the mass spectra for the HDX time course of two peptides in each of the three types of functional CF complexes, and compares the visual estimates of t_1/2_ (top) with the deconvolution method (bottom). For each HDX time course (vertical series of spectra) the spectrum shown in red and marked by # is the first one with ≥50% intensity in the high m/z mass envelope and/or the most symmetric pattern, indicating t_1/2_ for the EX1 exchange process has been reached. Each time course of spectra also includes a vertical green line at the m/z value halfway between unexchanged and back-exchanged, and a green asterisk marking the first spectrum in which the low m/z mass envelope has reached this halfway point. This identifies the t_1/2_ for the EX2 exchange process. These values are compared with the results of HX-Express, which deconvolutes bimodal patterns into two binomial distributions. The top table lists the intensity of the high m/z mass envelope, with the first value ≥ 50% highlighted in bold red indicating the t_1/2_ for the EX1 process. For the EX2 process, the bottom table lists the average mass and percent of exchange of the low m/z mass envelope, with the first value at ≥ 50% exchange highlighted in bold green, indicating the t_1/2_ of the EX2 exchange. The results of the deconvolutions demonstrate that the estimated t_1/2_ using our visualization method (Figure S3 top) is in good agreement with the HX-Express deconvolution results (Figure S3 bottom).

Visual estimates of t_1/2_ for EX1 and EX2 were performed for all peptides that exhibit a significant widening of the isotopic distribution during the HDX time course. Using measurements of peak width at 20% of max intensity, it has been previously demonstrated that a widening of > 2 Da during the time-course of HDX is indicative of an underlying EX1 process (20). The resulting estimated values of t_½_ for EX1 and EX2 exchange are shown in Figure 3 for functional complexes of CF4Q with CheA and CheW. The estimated values of t_1/2_ for all CF peptides are represented in colors ranging from dark maroon for short t_1/2_ (fast HDX) to pale purple for long t_1/2_ (slow HDX). When the first time point (3 min) exhibits complete exchange, this could be either an EX1 or EX2 process. Dark gray indicates peptides with very fast exchange (>70% exchange within 3 min). Light gray indicates peptides that display no evidence of EX1 exchange (≤ 2 Da widening of isotopic distribution), and black indicates regions with no peptide coverage. An average t_1/2_ estimate was calculated for overlapping peptides (black boxes in Figure 3A) and represented with the corresponding color on the CF monomer structures in Fig 3B to compare the EX1 (left) and EX2 (right) exchange profiles of CF4Q. For the 369-378 region, we considered the fact that EX1 exchange is more difficult to resolve for short peptides (eg. HDX for 2 peptides of 4-5 amino acids appears to be uncorrelated), and thus we used the t_½_ estimate of ≥ 960 min consistently observed for the three longer peptides (6-9 amino acids) in this region.

**Figure 3.**
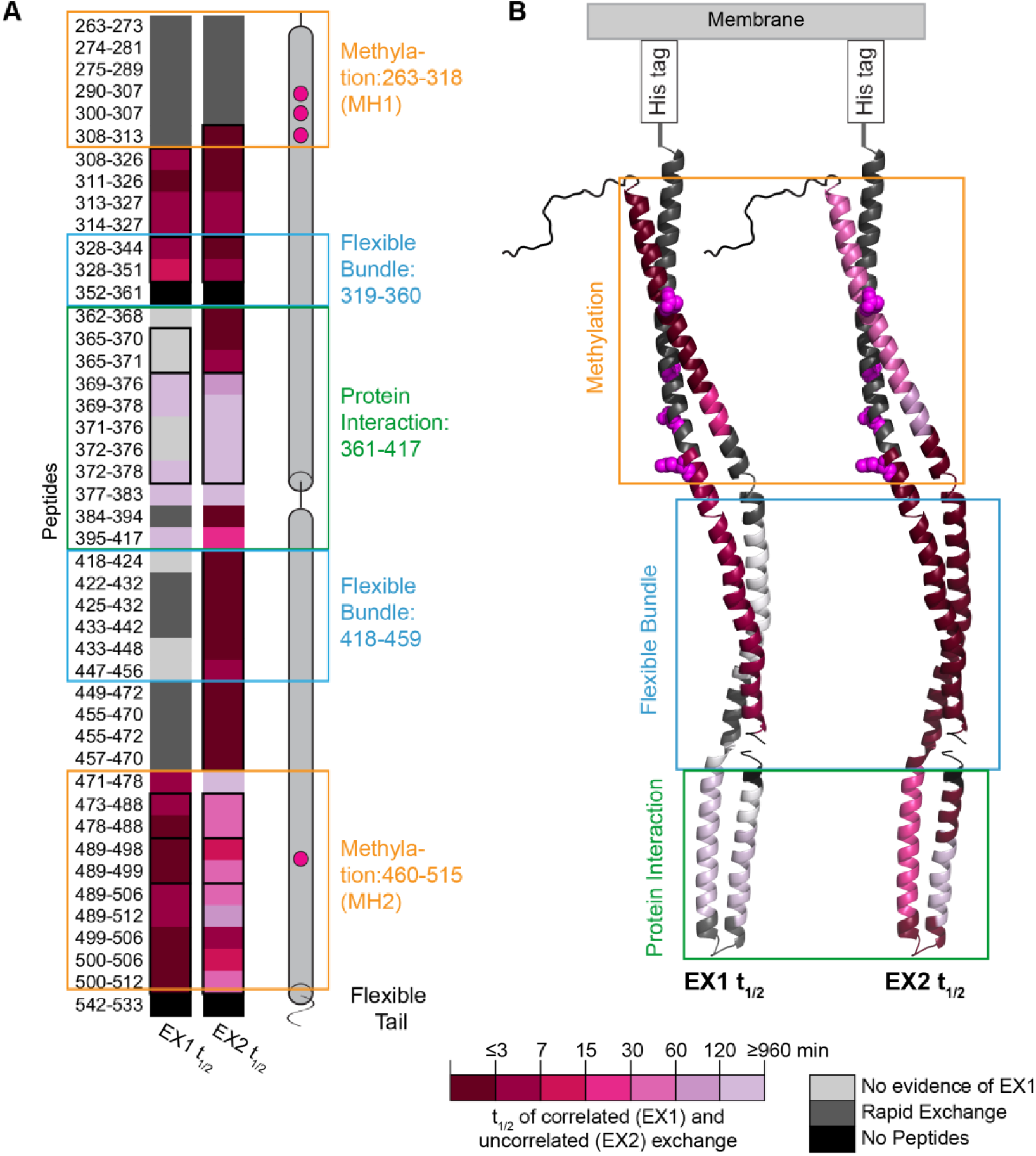
Estimated t_1/2_ of correlated (EX1) and uncorrelated (EX2) exchange of peptides throughout CF4Q in functional complexes with CheA and CheW. (A) The t_1/2_ values for EX1 and EX2 are represented by a color scale, from dark maroon to pale purple for fast to slow HDX (short to long t_½_). Dark gray represents peptides with > 70% exchanged in 3 min, which makes it either impossible to determine whether exchange is EX1 or EX2 (if HDX is complete in 3 min) or difficult to detect EX1 HDX. Light gray represents peptides with no detectable EX1 exchange. (B) CF monomer structure is colored according to the estimated EX1 t_1/2_ (left) and the estimated EX2 t_1/2_ (right) for CF4Q. Orange, green, and blue boxes outline the peptides and corresponding structure of the methylation, flexible bundle, and protein interaction subdomains of CF; sequence ranges for these have been deduced from comparative genomic analysis (21). The methylation sites (295, 302, 309, 491) are highlighted in magenta.

The emerging picture is that CF4Q within functional complexes exhibits widespread EX1 exchange. EX1 HDX processes are observed in all regions shown in maroon to pale purple, and may also occur in regions shown in gray. This suggests the entire CF populates an open state. The protein interaction sites exhibit slow EX1 (pale purple in Figure 3, t_1/2_ ∼16 hours) or no evidence of EX1 (light gray), but nearly all the rest of the protein exhibits very fast EX1 (t_1/2_ ≤ 7 min) (dark maroon) or fast exchange that makes it difficult to detect EX1(dark gray). Like the deuterium uptake analysis discussed above, this suggests that interactions between receptors, CheA, and CheW stabilize portions of the protein interaction region, but the rest of the CF (methylation and flexible bundle regions) samples an open and/or dynamic structure.

Functional complexes of CF4E and CF4Q.A411V exhibit similar HDX patterns, with widespread EX1. As discussed in the following section, HDX uptake is the same or slightly faster in these complexes, indicating that these states also have a stabilized protein interaction region with the remainder of the CF rapidly sampling an open structure.

Finally, the methylation region exhibits interesting asymmetry in its HDX behavior. Figure 3 shows that for segments where both EX1 and EX2 are detected, half times are similar in the protein interaction region (both slow) and flexible bundle region (both fast). Furthermore, both Figure 2B and Figure 3B show that the protein interaction and flexible bundle regions exhibit similarly slow or fast uptake across both the N and C terminal helices of the CF structure. In contrast, the methylation region exhibits much faster HDX on the N terminal side (MH1 HDX too fast to resolve) than on the C terminal side (MH2). Furthermore, MH2 is the only region in which the EX1 and EX2 kinetics are significantly different, with EX1 half times of 3-7 min and EX2 half times of 30-60 min. The MH2 helix is likely to be involved in stable packing interactions to slow its EX2 exchange, suggesting that in the methylation region MH2 forms the dimer interface and MH1 is highly dynamic.

### Inputs that favor the kinase-off state increase HDX rates throughout CF in functional complexes

To represent the kinase-off state we chose the A411V mutation, which has been shown to create a locked kinase-off receptor (13) and also to enhance methylation (14), as expected for the kinase-off state. This suggests it faithfully mimics the signaling state and is not just a disruption of kinase activation. Although A411 occurs near the top of the interaction region between CF and the P3 domain of CheA in the structural model, Figure 1 demonstrates that under our assembly conditions with excess CheA and CheW, this kinase-off mutant assembles complexes with native stoichiometry of ∼6 CF: 1 CheA: 2 CheW.

The differences in the deuterium uptake time course of the kinase-on (CF4Q) and kinase-off (CF4Q.A411V) functional complexes are illustrated in Figure 4 (see Figure S4 for complete set of peptides; colors in Figure 4 were chosen by averaging data in black boxes for overlapping peptides). Regions with no significant difference in uptake are represented in gray, based on a threshold for significance calculated from the standard deviations for the replicates for each state, as described in the Experimental Procedures. All significant differences are in the same direction: several peptides exhibit slower HDX in the kinase-on state (blue colors).

**Figure 4.**
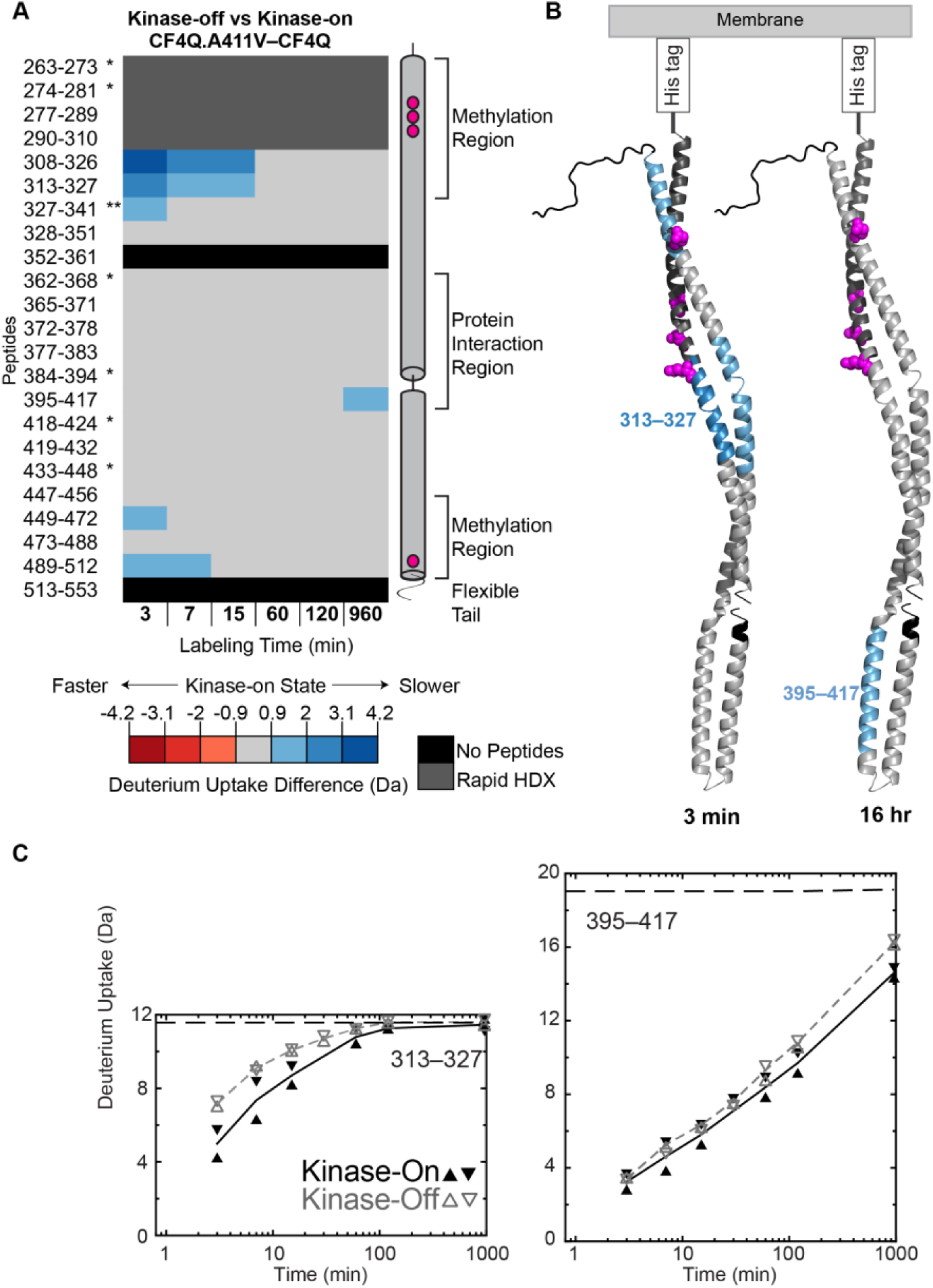
Difference in deuterium uptake between kinase-on and kinase-off signaling states. (A) Deuterium uptake of kinase-off CF4Q.A411V minus kinase-on CF4Q, for representative CF peptides covering the entire sequence, except segments shown in black. Segments in dark gray undergo very fast exchange (≥ 90% in 3 min), so these HDX experiments do not have the time resolution to detect differences between the two states in these regions. Segments in blue colors undergo slower HDX in the kinase-on state; segments in light gray show no significant difference in uptake (difference ≤ ±0.9 Da). Uptake differences are computed from averages of 2 replicates for each state, except for *peptides that were missing in one data set, and ** peptides that were missing in 2 of the 4 data sets. (B) CF monomer structure is colored according to the difference in deuterium uptake between the kinase-on and kinase-off states at the first (3 min) and last (16 hr) time points. (C) Plots of deuterium uptake vs time for representative peptides of both CF4Q (black symbols and solid lines) and CF4Q.A411V (gray symbols and dashed lines). Data points for all replicates are shown, along with lines connecting the averages of the replicates for each sample type. Dashed horizontal lines near top of plots indicate the deuterium level expected for complete exchange, based on control samples of CF4Q subjected to reversible heat denaturation in D_2_O buffer followed by the quench and mass spectrometry protocol used for all HDX samples.

Most peptides in and adjacent to the methylation region exhibit slower initial exchange in the kinase-on state, as shown by blue colors for early time points in Figure 4A and blue color in the top half of the structure in Figure 4B. Figure 4C includes the uptake plot for one of these peptides, 313-327: after an initial difference in uptake, the two curves converge at long times as the data approach complete exchange (dashed line is uptake level of completely-exchanged control). This suggests that the methylation region is less open/dynamic in the kinase-on state. Note that MH1 exchanges too quickly for detection of differences in HDX (exchange complete in 3 min), but a recent NMR study (14) of this segment within similar functional complexes indicates that the kinase-on state is less dynamic. There is also one region (473-488) of MH2 that exhibits no significant differences in HDX. With the exception of these regions, the entire top half of the CF exhibits less uptake at early times in the kinase-on state. This observation is consistent with our earlier study on similar samples of functional complexes (11).

At the 16 hr time point, a single peptide in the protein interaction region shows a significant difference in exchange. Peptide 395–417, on the C-terminal side of the hairpin tip, shows incomplete exchange in both CF4Q (4.4 Da) and CF4Q.A411V (2.8 Da). These data indicate significantly less deuterium uptake (1.6 Da) in the CF4Q kinase-on state, as represented by the blue color in Fig 4A-B and the difference in the uptake plot at long times in Figure 4C. This peptide, at the protein interface between CF and CheA-P3 (see Fig 2D), also contains the A411V mutation that creates the kinase-off state. We suggest that that the kinase-off state has a decreased affinity for CheA-P3, which leads to more complete HDX at long times. Such a decrease in affinity does not alter the amount of CheA incorporated and retained in the complexes (Table S1), perhaps due to the excess CheA/CheW conditions used and the high stability of the overall CheA/CheW hexagonal network of the array. Interestingly, the other peptides of the protein interaction region that show incomplete exchange at long times (N-terminal side of the hairpin tip: 369-383, see Figure S4) show no significant differences in uptake between the kinase-on and CF4Q.A411V kinase-off states.

Finally, although the HDX differences between the kinase-on CF4Q and kinase-off CF4Q.A411V are subtle, they demonstrate long-range coupling: differences are observed in both the methylation region at early times and in the protein interaction region with the kinase-off mutation site at long times. This is consistent with the fact that the A411V mutation causes both inactivation of the kinase (a local change in the CF-CheA interaction) and activation of methylation (a long-range change in the CF), and suggests that both of these activity changes are correlated with increasing the open/dynamic nature of the CF.

In contrast to the subtle differences in HDX between the kinase-on and kinase-off states, methylation has a much greater effect on the CF properties. Figure 5 illustrates the difference in deuterium uptake between unmethylated (CF4E) and methylated-mimic (CF4Q) functional complexes assembled with CheA and CheW on vesicles. Again, all significant differences are in the same direction, with peptides throughout CF exhibiting slower HDX in the methylated-mimic CF4Q (blue colors).

**Figure 5.**
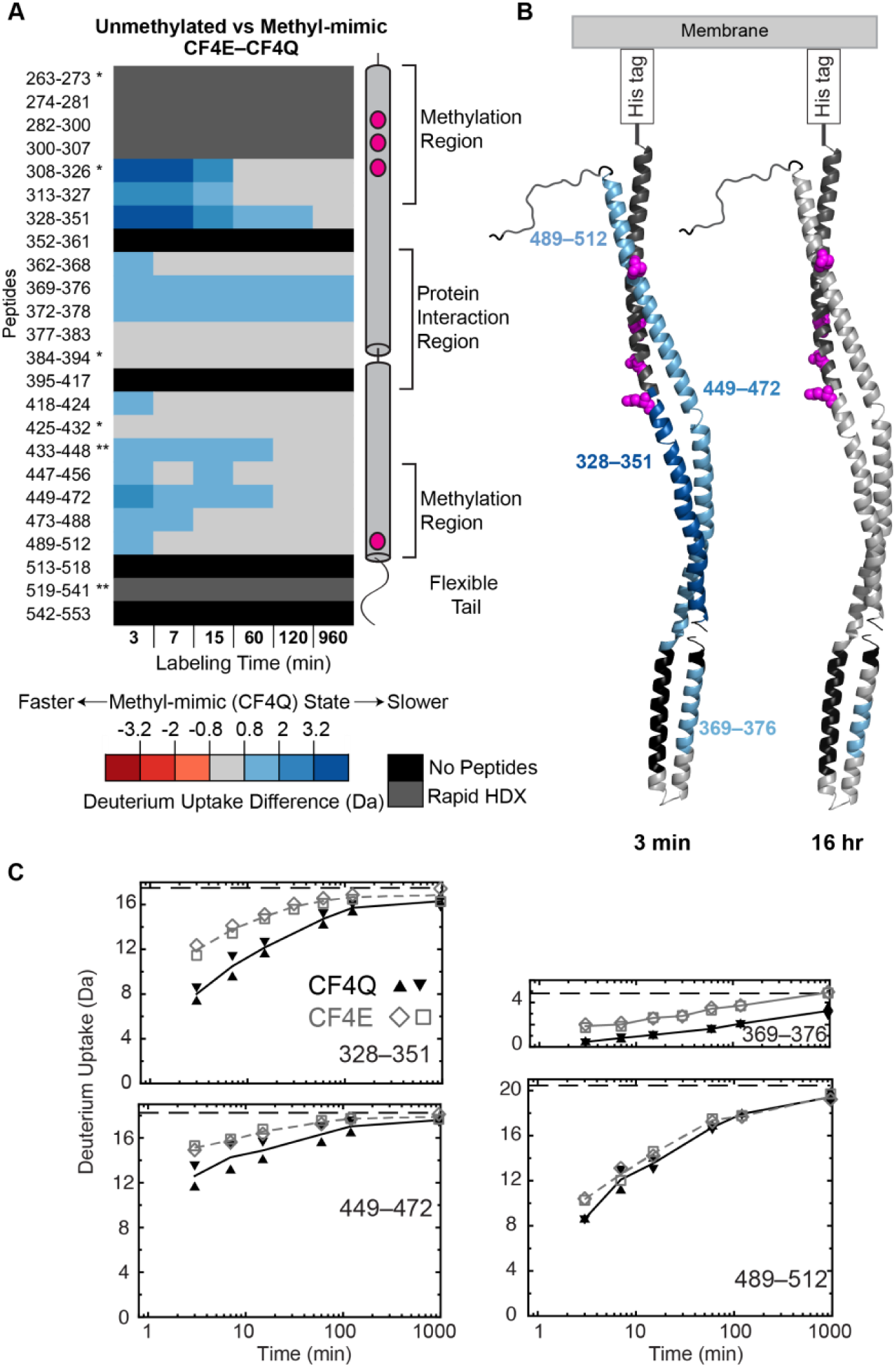
Difference in deuterium uptake between unmethylated and methylated-mimic states. (A) Deuterium uptake of unmethylated CF4E minus methylated-mimic CF4Q for representative CF peptides covering the entire sequence, except segments shown in black, with colors and replicates as described for Figure 4. Segments in blue colors undergo slower HDX in the methylated-mimic state. (B) CF monomer structure is colored according to the difference in deuterium uptake between the unmethylated and methylated-mimic states at the first (3 min) and last (16 hr) time points. (C) Plots of deuterium uptake vs time for representative peptides of both CF4Q (black symbols and solid lines) and CF4E (gray symbols and dashed lines). Data points for all replicates are shown, along with lines connecting the averages of the replicates for each sample type. Dashed horizontal lines near top of plots indicate the deuterium level expected for complete exchange, based on control samples of CF subjected to reversible heat denaturation in D_2_O buffer followed by the quench and mass spectrometry protocol used for all HDX samples.

At early times, peptides throughout the methylation, flexible bundle, and protein interaction regions exhibit slower exchange in the CF4Q methylated-mimic state (Fig 5A, B & C). In fact, of the regions with measurable HDX rates, nearly all contain at least one peptide that shows significantly less deuterium uptake in CF4Q (see Figure S4). The single exception is the 379-394 region, which is mostly the hairpin tip, 384-394, that exhibits fairly rapid exchange in all complexes. Thus, we conclude that methylation causes widespread stabilization (resulting in slower initial HDX) throughout all regions of the CF with measurable HDX rates (all except the hairpin tip and N and C-terminal regions).

At long times (16 hr), peptides involved in protein interactions (those that show incomplete exchange) exhibit reduced uptake in the methylated-mimic state. This is observed on the N-terminal side of the hairpin tip (369-378, which interacts with CheA-P5/CheW/CF as shown in Figure 2D). Unfortunately, the peptide involved in protein interactions on the C-terminal side (395-417) is not detected in the CF4E data sets, so we cannot determine whether it too would show decreased uptake at long times in CF4Q relative to CF4E. Although it is not clear whether methylation alters CF interactions with CheA-P3, it appears that methylation increases the affinity of CF interactions with CheA-P5 and/or CheW and/or other CF dimers in the trimer of dimers. Overall, these HDX data suggest that methylation causes widespread stabilization of CF and its interactions.

Although the location and relative magnitude of the individual HDX differences between states are interesting, the key observation is that all of the differences are in the same direction, pointing to a destabilization throughout the CF by conditions that favor the kinase-off state (demethylation and locked-off mutant).

### Signaling states primarily alter the fraction of the fast phase of HDX

Uptake plots show that HDX of most peptides is biphasic, with an initial fast phase that is often too fast to quantify in our experiments (first time point at 3 minutes) followed by a slow phase. Figure S5 shows uptake plots for representative peptides throughout the CF. These data were fit to obtain the fraction and rate constant of fast uptake (f_1_, k_1_) and slow uptake (f_2_, k_2_), which are tabulated in Table S2. For the many cases of very fast uptake that cannot be defined by our data, Figure S5 shows that a faster uptake (dotted curves with larger k_1_) fits equally well, so a lower limit for k_1_ is reported in Table S2. The fastest uptake (monophasic, very fast) is exhibited by MH1 (260-273) and the C terminal region (526-532). The membrane-distal tip of the receptor (384-394) also exhibits predominantly (≈90%) very fast uptake (k_1_ ≥ 0.7 min^-1^), with a minor slow phase (k_2_ ≤ 0.04 min^-1^). The slowest uptake, observed in most of the protein interaction region, is biphasic: with the exception of 362-368, the protein interaction region peptides of CF4Q exhibit ∼1/3 or no fast phase (k_1_ ≈ 0.3 min^-1^) and 2/3 or all slow phase (k_2_ = 0.006 min^-1^). The rest of the CF exhibits biphasic uptake, with approximately 2/3 very fast to fast uptake (k_1_ ≥ 0.4 min^-1^, except all fast for 455-470) and 1/3 slow uptake (k_2_ < 0.04 min^-1^, except for 422-432). These values for 4Q, along with those for 4E and 4Q.A411V are listed in Table S2.

The uptake plots for selected peptides (Figure 4C and 5C) and for representative peptides throughout the CF (Figure S5) show that the uptake differences between CF4Q, CF4E, and CF4Q.A411V are primarily due to a change in the rapid initial phase of HDX. Comparison of the best fit parameters listed in Table S2 confirms that changes between states are primarily an increase in the fraction of the fast phase in CF4E (f_1_ in bold in Table S2): the average fraction of this phase increases from 0.7 in CF4Q to 0.8 in CF4E.

Superpositions of the mass spectra of CF4Q and CF4E arrays after 3 min of HDX were used to determine whether the change in fast uptake is a change in the EX1 or EX2 HDX. Figure S6 shows superpositions for peptides throughout CF that show both a change in uptake and resolved EX1 at 3 min. Some of these peptides exhibit faster initial EX2 in CF4E, as indicated by a horizontal shift (blue arrow) in the tallest peak of the low m/z distribution. All of these peptides exhibit faster initial EX1 in CF4E, as indicated by red vertical arrows showing the decrease in the area of the low m/z distribution and the increase in the area of the high m/z distribution. Although CF4E exhibits slightly faster EX2 in a few peptides (314-327, 328-351, 433-448), the changes in EX1 are larger. Therefore the primary observed change is that initial EX1 uptake is faster in CF4E than in CF4Q.

Although it was not possible to deconvolute all of the mass spectra to obtain EX1 and EX2 uptake plots, HXpress deconvolutions were successful for a series of peptides covering the MH2 region (471-512). These mass spectra could be deconvoluted because the EX1 uptake occurs more quickly than EX2 uptake, so the two isotopic distributions remain resolved through most of the time course, until the EX2 envelope catches up to the fully exchanged envelope. It is clear from the estimated t_1/2_ values shown in Figure 3 that this difference in rate, with EX1 >> EX2, occurs only in MH2. The deconvoluted uptake data plotted in Figure 6 shows that the EX1 uptake (dashed lines) is biphasic, with a large initial fast phase that occurs during the dead time of the measurement, followed by a slow phase monitored by the experiment. CF4E (blue) exhibits an increase in the fast EX1 and no change in the rate of slow EX1. EX2 (solid lines) has little or no unresolved fast phase, followed by slow uptake at a rate comparable to the EX1 slow uptake. There is no significant change in the EX2 uptake between samples. Thus HXpress deconvolutions of MH2 peptides (471-512) show that all changes in uptake are due to an increase in the magnitude of fast EX1 in 4E, consistent with the conclusions suggested by superpositions of spectra for peptides throughout CF after 3 min of HDX.

**Figure 6.**
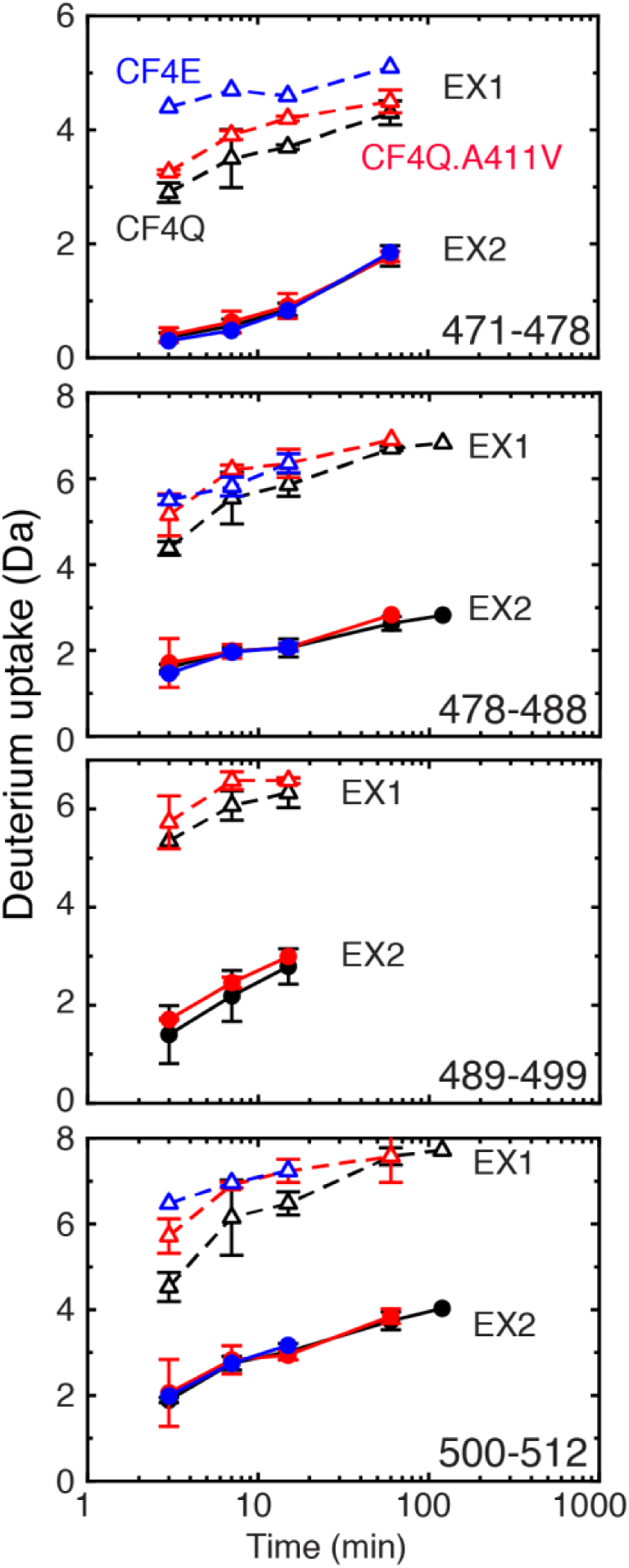
EX1 and EX2 uptake for four peptides from methylation helix 2 (MH2). The EX1 uptake (open triangles and dashed lines) for all three states has a rapid initial phase followed by a slower phase with comparable rate to the EX2 uptake (filled circles and solid lines). CF4Q uptake is plotted in black, CF4Q.A411V in red, and CF4E in blue. Deconvolutions were performed with HXpress.

## Discussion

### Diverse dynamic properties of CF in functional signaling complexes

This study is the first to measure hydrogen exchange of the CF in samples where all of CF is incorporated into functional complexes with CheA and CheW, providing new insights into its diverse dynamics and stability properties. Furthermore, the large number of short peptides (∼74 per state with an average length of 12) define the boundaries of regions with different HDX properties, which was not possible in our previous study (only 26 peptides with an average length of 20). The HDX uptake rates and patterns for CF lead to two significant new proposals: (1) the receptor cytoplasmic domain remains partially disordered within functional complexes, and (2) signaling occurs via modulation of this disorder/stability by both ligand binding and methylation.

HDX-MS data suggest that (1) regions near the N and C termini are unstructured (very fast HDX, complete in 3 min), (2) regions involved in protein interactions near the membrane-distal tip are stable (slow HDX, incomplete at 16 hr) and (3) the rest of the CF exhibits intermediate properties, including fast HDX which is ≥ 50% complete in 15 min. Interestingly, these diverse HDX behaviors are localized within the receptor regions previously identified by bioinformatic analysis (21): the protein interaction region (with the exception of the 384-394 tip) has slow HDX, the flexible bundle region has fast HDX, and the methylation region has asymmetric behavior, very fast HDX for MH1 and a combination of fast EX1 and intermediate EX2 for MH2. Thus HDX has identified distinct dynamic properties of previously identified functional regions of the receptor cytoplasmic domain within functional, membrane-bound signaling complexes with CheA and CheW.

The very fast HDX near the N and C termini is consistent with the results of other studies. The C terminal tail has been previously shown to be highly mobile by EPR (17), HDX (11), and NMR (14), and is thought to function as a flexible tether to the CheR and CheB binding site at the C terminus of the receptor (22). Near the N terminus, we observe very fast HDX throughout methylation helix 1 (MH1). This agrees with our previous HDX and NMR studies on similar samples, which found MH1 dynamics in both CF4E and CF4Q to be comparable to the C terminal tail dynamics (11, 14). EPR studies of site-specific spin labels measured the mobility of helices of the cytoplasmic domain of intact Asp receptor inserted into nanodiscs, but lacking CheA and CheW (16). Consistent with our results, Bartelli and Hazelbauer observed that MH1 is more dynamic than MH2. Based on EPR lineshapes at sites throughout the receptor cytoplasmic domain, they suggested the protein conformation samples a disordered helical backbone, with the greatest fraction of disorder observed in MH1 of the 4E receptor (16). The high mobility of MH1 is likely to be characteristic of the native receptor, since it has been observed in both nanodisc-imbedded intact receptor dimers that lack CheA and CheW (EPR study) and in membrane-bound complexes of CF, CheA, and CheW that lack the periplasmic and transmembrane receptor domains but display native array structure and function (this study).

The dual nature of HDX in the rest of the CF, with fast exchange everywhere except the sites of protein contacts, suggests that assembly of CF into functional complexes stabilizes it primarily in the protein interaction region. This is further supported by the observation that CF exhibits widespread EX1 exchange (see Fig 3), which indicates the protein is sampling an open state. The very slow EX1 observed in the protein interaction region suggests that in this region the CF samples the open state when the interacting proteins dissociate, which is likely slow due to the network of multiple interactions in the hexagonal array. The CF exhibits several indicators of a dynamic/open structure. (1) Rapid EX1 (short t_½_ values except the protein interaction region) indicates rapid unfolding/opening. (2) Rapid EX2 (short t_½_ values except in MH2) indicates low stability. (3) Biphasic uptake with 66% fast HDX (on average for CF4Q, outside of the protein interaction and very fast exchange regions) suggests an unstable fraction. The latter suggestion is based on a comparison of protection factors (PF) for these phases (Table 1) to those measured for ACBP, another four-helix bundle protein (23). For the slow uptake phase, log(PF) > 4 for most peptides, comparable to protection factors of alpha helices in ACBP (23).

The fast uptake phase exhibits much lower protection factors, with an average log(PF) =2.9, suggesting this is an unstable fraction. This fast phase fraction cannot be denatured protein because (1) the high activity of these complexes is inconsistent with 2/3 being denatured, and (2) the fast phase fraction is not uniform throughout the CF, with only 1/3 fast phase in most of the protein interaction region. Also, the fast uptake fraction cannot be in rapid equilibrium with the 34% slow uptake fraction or the entire uptake would be very fast.

A possible explanation of the biphasic HDX behavior is the structural inequivalence of the receptors within functional arrays. The three dimers of each trimer of dimers make different protein contacts within the hexagonal array with CheA and CheW (discussed below). We suggest that all CF dimers are stabilized by this network of contacts in the protein interaction region, but beyond this region perhaps only one dimer in each trimer of dimers in the array is stabilized and accounts for the 34% slow exchanging fraction. Electron cryotomography of arrays of intact receptors (1, 2) and receptor fragments (24) have both shown that the baseplate region of the receptor interacting with CheA and CheW has well-defined electron density, but the electron density becomes less well-defined in the membrane-proximal regions. This may reflect the dynamics/instability that we report, but in our samples these dynamics begin just above the protein interaction region in about 2/3 of the CF’s.

Our data also provide new insights into the properties of the methylation region. Methylation helix 2 (MH2) peptides exhibit about 2/3 fraction with fast uptake, which is predominantly EX1. This suggests that MH2 in two of the three dimers undergoes rapid unfolding/opening to an open state (rapid EX1). The slower EX2 in this region, with log(PF) > 4, suggests that in the third dimer MH2 must pack against another helix. This suggests that MH2/MH2’ forms the dimer interface, since MH1 shows no evidence of stable packing (log(PF) ≤ 2.4). Such a model is consistent with a recent mutagenesis study suggesting stable MH2/MH2’ packing at the membrane proximal end of MH2 in the Ser receptor (25), but is difficult, however, to reconcile with the existence of disulfide crosslinked MH1 /MH1’ dimers within this region that retain activity (5, 26).

Identification of peptides with incomplete HDX at 16 hr also provides experimental evidence for the protein-protein interactions present in functional chemoreceptor arrays. Receptor interface residues identified by chemical shift perturbation in solution NMR experiments on binary complexes of protein fragments (27, 28) or disulfide crosslinking of intact proteins (29, 30) are listed in Table S3. Most of these residues are found within the peptides that show incomplete HDX at 16 hours. The two exceptions (385 and 386, highlighted in bold red in Table S3) are found within the receptor tip peptide that shows rapid HDX (384-394). These differences may be due to the longer reach of the disulfide crosslinking approach (4.6 Å between beta carbons (29)) or to conformational changes induced in the receptor tip upon protein binding. Note that the contacts identified by NMR using a P4-P5 construct of CheA (27) all lie within 372-383, consistent with our suggestion that incomplete HDX of this peptide is due to interactions with P5. Furthermore, this indicates that incomplete HDX of the 395-417 peptide is likely to be due to receptor/P3 interactions that are not present in the NMR experiment. Finally, Table S3 shows good agreement between the HDX results and the contacts (receptor residues within 3 Å of CheA or CheW) observed in two structural models derived from x-ray crystallography, electron cryotomography, and molecular modeling (1, 31).

### Proposed signaling mechanism: long-range allostery via stabilization

A primary goal of our study was to compare the dynamics of the receptor in different signaling states and methylation states, both to test the proposed inverse dynamics model and to gain further insights into the signaling mechanism. The HDX results indicate that *both* the methylation region and protein interaction region of CF incorporated in functional complexes are stabilized in the kinase-on state relative to the kinase-off state (Figure 4), and in the methylated state relative to the demethylated state (Figure 5). This is inconsistent with one part of the inverse dynamics model (7) that proposed the methylation region is stabilized but the protein interaction region is *destabilized* in the kinase-on state. Decreased HDX in both regions was also observed in our previous HDX study (11) on arrays of CF4E assembled with CheA and CheW at high density (kinase-on state) relative to those assembled at low density (kinase-off state). An EPR study of intact Asp receptors in nanodiscs similarly observed decreased dynamics in the methylation region for the methylated state relative to the demethylated state (16). Finally, EPR/DEER studies of constructs containing the Asp receptor methylation and protein interaction domains linked to variants of the Aer HAMP domain observed decreased protein interaction region dynamics in the kinase-on constructs relative to the kinase-off constructs (32). Results of all of these studies using different methods and samples indicate that inputs favoring the kinase-on state lead to reduced dynamics in both the methylation region and the protein interaction region.

Based on our HDX results we propose a new model for the signaling mechanism, illustrated in Figure 7: we propose that long-range allosteric signaling through the cytoplasmic domain occurs via stabilization of its open/dynamic structure. In the absence of CheA and CheW, the cytoplasmic domain is known to be highly dynamic. For example, this domain of the intact Asp receptor exhibits promiscuous disulfide crosslinking (5) and the Asp receptor CF in solution exhibits properties similar to a “molten globule,” with an alpha helical secondary structure but lacking a well-defined tertiary structure (ANS fluorescence due to exposed hydrophobic groups, rapid proteolysis, and very fast amide hydrogen exchange (4)). As discussed above, we propose that binding of CheA and CheW primarily stabilizes the protein interaction region (yellow and orange in Figure 7). This binding also partially stabilizes the rest of the CF (most of which no longer has very fast HDX), but it retains some open/dynamic “molten globule” characteristics, such as fast HDX and EX1.

**Figure 7.**
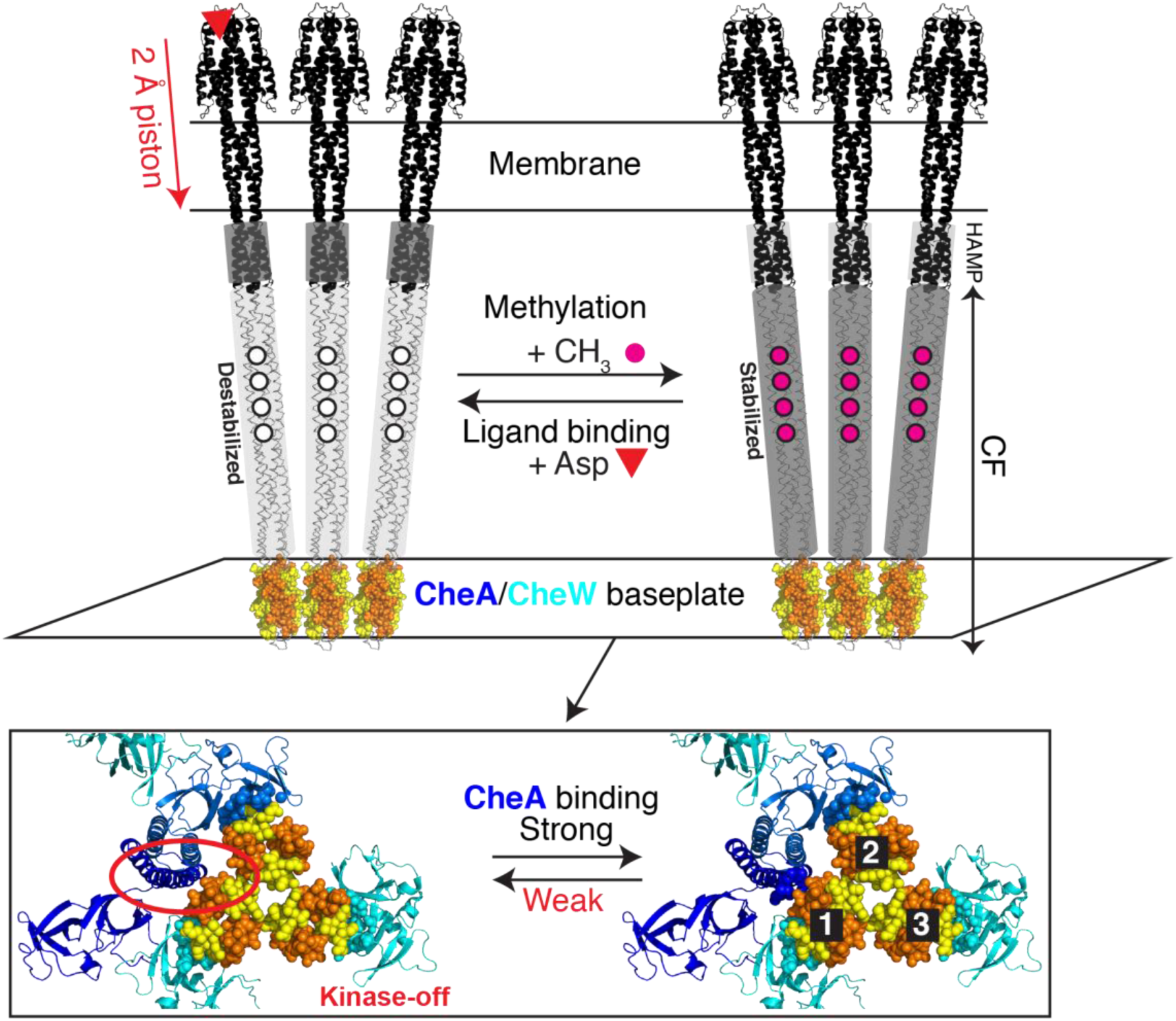
Proposed long-range allostery via protein stabilization. Top: Model of the receptor trimer of dimers in both signaling states. The cytoplasmic domain consists of the elongated CF connected by the small, membrane-proximal HAMP domain to the periplasmic and transmembrane domains. HDX data indicate that interactions with CheA and CheW in the baseplate of the hexagonal array stabilize only this region of the CF (yellow/orange). The remainder of the CF (backbone trace) exhibits widespread EX1 and fast or very fast exchange. Although we do not have HDX data for the HAMP domain, it is represented in the opposite color (dark vs light gray) based on the proposal that the methylation region and HAMP have inverse stability (6). Bottom: view from the membrane of the trimer of dimers protein interaction region. Interactions with the CF (yellow and orange regions with incomplete HDX as shown in Figure 2) are highlighted using spherical representation for residues of CheA (blue) and CheW (cyan) within 10 Å of CF, illustrating the inequivalent interactions made by the three CF dimers. We propose that CheA binds to CF in a kinase-on conformation and stabilizes CF (dark gray, right). Thus anything that destabilizes CF (light gray), such as the ligand-induced piston or demethylation, weakens CheA binding. We suggest that loss of a key interaction (eg interactions between CF dimer 1 and the P3 domain, enclosed by the red oval), causes CheA to revert to the kinase-off state. Note that CheA would not dissociate completely from the arrays due to the network of other interactions.

We propose that signaling inputs control the kinase activity of CheA by modulating the stability of the cytoplasmic domain, which in turn modulates interactions with CheA. CheA in solution has very low kinase activity. Binding of CheA and CheW stabilizes CF, and CheA binds in a kinase-on conformation. Since both a kinase-off mutant (CF4Q.A411V) and the unmethylated state (CF4E) exhibit faster HDX, primarily due to an increase in the fast EX1 exchange, we suggest that the ligand-induced piston and demethylation each destabilize the CF. This is represented in Figure 7 by the dark gray (stabilized) kinase-on state CF (right) becoming light gray (destabilized) in the kinase-off state (left). Such destabilization of CF in turn disfavors the binding interactions with CheA and CheW. CheA and CheW do not dissociate from the array due to the network of interactions among the proteins in the array. But we propose that CheA loses a key contact with the receptor and reverts to its kinase-off state. Thus both CheA binding and signaling inputs modulate stability of CF. This model is consistent with the observed parallel changes in HDX in the methylation and protein interaction regions of the kinase-off state: both regions are destabilized by inputs that favor the kinase-off state.

The CheA/CheW binding interactions with CF and potential signaling-related changes are illustrated in the structural model of the baseplate of the array (viewed from the membrane) at the bottom of Figure 7. This figure shows only the protein interaction regions of CF that exhibit incomplete HDX at long times, colored yellow and orange as in Figure 2. To highlight the inequivalent protein interactions of CF dimers within the array, residues of CheA and CheW within 10 Å of CF are shown as spheres (right side). Dimers 1 and 2 interact with the CheA/CheW ring in the array, with dimer 1 interacting via yellow residues 372-383 with CheW and via orange residues 395-417 with P3 of one CheA (darker blue). In the other dimers, yellow residues 372-383 mediate different interactions: dimer 2 interacts with P5 of the other CheA (lighter blue) and dimer 3 interacts with CheW of a W-only ring.

One specific proposal consistent with the HDX data and current structural models is that upon destabilization by ligand binding or demethylation CF loses a key contact with CheA-P3, which causes CheA to revert to the kinase-off state. This is based on the higher total HDX observed for the 395-417 peptide of CF4Q.A411V (a peptide that is unfortunately not observed in CF4E), which suggests that it is the interaction with CheA-P3 that is weakened in the kinase-off state. It is possible that the A411V mutation (contained in this peptide) weakens the receptor-CheA interaction, but the remaining network of interactions are sufficient for assembly of arrays *in vivo* (13) or *in vitro* under excess CheA conditions (this study). In contrast, the peptides involved in interactions with CheA-P5 and CheW (372-383) show changes in fast phase HDX in CF4E (which is kinase-on under our assembly conditions) but not in CF4Q.A411V, suggesting that these are not the key contacts that control the kinase. This proposal is also consistent with observations that ligand binding changes disulfide crosslinking rates between CheA and receptor residues 396 and 399, although the CheA crosslinking partners were in P5 rather than in P3 (29). The proposed loss of P3/CF contact(s) is illustrated in Figure 7 (red oval on the left) by depicting P3 contact residues without spheres, to represent loss of the interaction with CF residues 395-417. Within the current structural model, this would yield a state with the receptor trimer of dimers contacting only one monomer of the CheA dimer, as shown for the kinase-off state on the left in Figure 7, which may be insufficient to position the CheA domains for productive trans-phosphorylation across the dimer interface.

In summary, HDX suggests the following key elements of the proposed model for long-range allosteric coupling via protein stabilization. (1) The cytoplasmic domain is intrinsically unstable even within functional complexes; it is stabilized in complexes only in the small protein interaction region, and the asymmetry of these contacts leads to differential stabilization of different dimers in the rest of the CF. (2) Signaling inputs destabilize the cytoplasmic domain (below HAMP), which weakens binding interactions and causes loss of key contacts with CheA to turn off its kinase activity. Our HDX data also suggest that the key contacts controlling kinase activity are with P3.

A key challenge for testing this and other models for the signaling mechanism is the inequivalence of receptor interactions within functional receptor arrays. Arrays contain three types of receptor dimers: dimer one contacts CheA-P3 and CheW, dimer two contacts CheA-P5, and dimer three contacts CheW. This inequivalence makes properties and signaling-related changes of each type of dimer more difficult to observe, due to the background of the other two types of dimers. For instance, if the kinase-off A411V mutation exerts its effects by destabilizang interactions with CheA P3, this would alter only one of the six CF in each trimer of dimers. In contrast, the 4E methylation state would alter the stability of all six CF. This difference may account for the observation of greater HDX changes in 4E than in A411V.

We propose a signaling mechanism in which both protein-protein interactions and signaling inputs modulate stability of an open/dynamic domain of a receptor. For a system like the chemoreceptor with frequent on/off switching, such a mechanism would be inefficient if it involved complete dissociation of the binding partner. Thus one potential function of the chemoreceptor array is to position other proteins (eg CheW in the CheA/CheW rings) that maintain CheA in close proximity to the receptor, regardless of the receptor stability, for efficient gain/loss of a key contact that controls the kinase activity. By this reasoning, such allosteric coupling via binding-modulated stabilization may also be found in other multi-protein complexes where interactions with other proteins position a receptor binding partner such that receptor destabilization could break key binding interactions without complete dissociation of the binding partner from the complex.

Modulation of a coupled equilibrium between CheA binding and stabilization of a disordered CF is an exciting new idea for explaining the integration of different inputs (ligand-induced piston and methylation) and the long-distance propagation of the signal 200Å from the membrane to the membrane-distal-tip of the chemotaxis receptor. Interestingly, in the related sensor histidine kinases (which contain the sensor and kinase within the same protein), signaling-related local unfolding of helices has been observed by HDX-MS (33, 34). Additional study of the role of folding/stabilization in chemoreceptor signaling is important to provide a detailed understanding of the mechanism and how it compares to observations in other systems in which more localized folding/unfolding is key to signaling mechanisms (35), and in cases where intrinsically disordered domains mediate allosteric interactions (36).

## Experimental Procedures

### Protein Expression and Purification

#### His-tagged Asp receptor cytoplasmic fragment

(CF) begins with an N-terminal His tag, followed by the last 5 residues of the HAMP domain and then the methylation region. CF was expressed from plasmids pHTCF4Q, pHTCF4E (8) or pCF4Q.A411V (37), all of which encode ampicillin resistance. BL21(DE3) *E. coli* cells (or DH5□F’ for CF4E) co-transformed with one of these plasmids (amp^R^) and with pCF430 (tet^R^), were streaked on Luria-Bertani (LB) agar plates with ampicillin (150 µg/mL) and tetracycline (10 µg/mL) and incubated at 37°C. For each CF, single colony was inoculated into 2 mL LB broth containing the same concentration of antibiotics, grown at 37°C with 200 rpm shaking until the optical density at 600nm (OD_600_) reached ∼0.6, and then used to inoculate a 2 L culture. This large scale culture was incubated in a 37°C shaker until OD_600_ reached ∼0.6, induced with 1mM IPTG, and grown overnight at 10°C. The cells were harvested by centrifugation at 3500 rpm (Beckman Coulter Allegra 6R Tabletop centrifuge, G38 swinging bucket rotor) at 4°C for 30 min, and then resuspended in lysis buffer (75 mM K_x_H_x_PO_4_, 500 mM NaCl, 5 mM imidazole, 1mM EDTA, pH 7.5) and lysed using a microfluidizer (Microfluidics, Inc) at 16K psi on ice. PMSF (1 mM) was added every hour to prevent proteolysis. Cell debris was separated using centrifugation at 10,000 rpm at 4°C for 90 min in a Sorvall RC-5B centrifuge, SS-34 rotor. The supernatant was then passed through a HisTrap FF Ni^2+^-NTA affinity column (GE Healthcare) that had been equilibrated with 10 column volumes of equilibration buffer (75 mM K_x_H_x_PO_4_, 500 mM NaCl, 5 mM imidazole, pH 7.5). The column was washed with 10 column volumes of wash buffer (75 mM K_x_H_x_PO_4_, 500 mM NaCl, 50 mM imidazole, pH 7.5) to remove loosely bound proteins. CF was eluted with elution buffer (75 mM K_x_H_x_PO_4_, 500 mM NaCl, 500 mM imidazole, pH 7.5) and located in the fractions by SDS-PAGE. Fractions containing CF were combined and treated with 5 mM EDTA to chelate any Ni^2+^ that came from the column. The CF solution was then dialyzed at 4°C against dialysis buffer (75 mM K_x_H_x_PO_4,_ 75 mM KCl, pH 7.5) in 7 kDa molecular weight cutoff dialysis tubing. Typically the dialysis was against 2L of dialysis buffer overnight, and then an additional 4 hour dialysis against 2L of fresh dialysis buffer, so that the calculated remaining EDTA and imidazole concentrations would be less than 0.01mM. The concentration of CF was determined using a BCA assay (Thermo Scientific).

#### TEV-cleavable His-tagged CheA and CheY

were expressed using plasmids pTEV-cheA (kan^R^) and pTEV-cheY (kan^R^) (14) in BL21(DE3) with 50 µg/mL kanamycin. Protein expression and growth protocols were the same as for His-tagged CF except CheA was induced at 37°C for 3 hours, and CheY was induced overnight at 4°C. The same purification protocol was used to purify CheA and CheY, except using different buffers: lysis buffer (75 mM Tris-HCl, 100 mM KCl, 1 mM EDTA, pH 7.4), equilibration buffer (75 mM Tris-HCl, 100 mM KCl, pH 7.4), wash buffer (75 mM Tris-HCl, 100 mM KCl, 10 mM imidazole, pH 7.4), elution buffer (75 mM Tris-HCl, 100 mM KCl, 250 mM imidazole, pH 7.4), and dialysis buffer (75 mM Tris-HCl, 100 mM KCl at pH 7.4). Proteins were concentrated using 10 kDa cutoff Amicon filters centrifuged at 3000 rpm in a Beckman Coulter Allegra 6R Tabletop centrifuge, G38 swinging bucket rotor at 4°C until the volume was around 3 mL. Concentrations were estimated using the absorbance at 280 nm and extinction coefficients of 25000 M^-1^cm^-1^ for CheA and 6970 M^-1^cm^-1^ for CheY (38)) to choose conditions for TEV cleavage.

#### TEV-cleavable His-tagged CheW

was expressed using the plasmid pSJCW grown in DH5α *E. coli* cells, and protein purification was performed as described above except that after lysis the cell debris was spun down in a pre-chilled centrifuge (Sorvall, SS-34 rotor) at 10,000 rpm at 4°C for 1 hours followed by 1 hour of ultracentrifugation (Beckman Optima™ LE, 80K, SW28 rotor) at 28,000 rpm (104000 x g) at 4°C. Purification buffers were lysis buffer (50 mM K_x_H_x_PO_4_, 300 mM KCl, 10 mM imidazole, pH 8.0), wash buffer (50 mM K_x_H_x_PO_4_, 300 mM KCl, 50 mM imidazole, pH 8.0), elution buffer (50 mM K_x_H_x_PO_4_, 300 mM KCl, 250 mM imidazole, pH 8.0), and dialysis buffer (75 mM K_x_H_x_PO_4_, pH 7.5). CheW was concentrated as described above and its concentration was estimated using the absorbance at 280 nm and an extinction coefficient of 5120 M^-1^cm^-1^ (39).

#### TEV-protease

with an N-terminal His tag was expressed from the plasmid pRK793 (amp^R^), a gift of D. Waugh (Addgene plasmid 8827 (40)). BL21(DE3)-RIL (cam^R^) cells containing this plasmid were grown in cultures containing 150 µg/mL ampicillin and 50 µg/mL chloramphenicol at 37°C until OD_600_ reached 0.6–0.8, expression was induced with 1 mM IPTG for 4 hours, and TEV protease was purified as described above for purification of CF.

#### His-tag cleavage of CheA, CheW and CheY

was performed at a 50:1 ratio of His-tagged protein: TEV-protease. The mixture was stirred at 4°C overnight and an additional 2 hours at 25°C. Cleavage by TEV was assessed by SDS-PAGE gel shift; if incomplete cleavage was observed, the sample was again incubated overnight at 4°C and assessed by SDS-PAGE. Cleaved proteins were then purified using the HisTrap column as described above, except the TEV protease was bound to the column and the flow-through contained the TEV-cleaved proteins. Proteins were then concentrated to the desired concentrations using 10 kDa molecular weight cutoff Amicon filters as described above. BCA assay (Thermo Scientific) was used to determine the final concentrations of all CF, CheA, CheW and CheY solutions. Proteins were aliquoted into 1.5 mL Eppendorf tubes, flash frozen with liquid nitrogen, and stored at -80°C.

### Lipid Vesicles

Vesicles were prepare using DOPC (1,2-dioleoyl-*sn*-glycero-3-phosphocholine) and the nickel-chelating lipid DGS-NTA-Ni^2+^ (1,2-dioleoyl-*sn*-glycero-3-{[*N*-(5-amino-1-carboxypentyl)-iminodiacetic acid]succinyl}) (Avanti Polar Lipids) mixed in a 1.5:1 ratio, and vesicles were prepared as previously described in kinase buffer (50 mM K_x_H_x_PO_4_, 50 mM KCl, 5 mM MgCl_2_, pH 7.5) including five freeze/thaw cycles and extrusion with 15 passes through a 100 nm diameter pore size membrane (37). A total stock concentration of 3 mM lipid (DOPC and DGS-NTA-Ni^2+^) was used to assemble functional complexes; a final concentration of 725 µM lipid was used in the assembly. This lipid concentration was chosen to accommodate all of the 30µM CF in hexagonal arrays. Concentrations of 12 µM CheA and 24 µM CheW were chosen to yield maximal kinase activity and native stoichiometries of ∼6 CF:1 CheA: 2 CheW (41).

### Vesicle-mediated Assembly of Functional Complexes

Functional complexes (8) with native-like structure (9) were assembled by combining (in the following order) autoclaved water, kinase buffer, 1 mM PMSF in ethanol, 12 µM CheA, 24 µM CheW, 30 µM CF (CF4Q, CF4Q.A411V or CF4E), and 725 µM vesicles. Complexes were incubated overnight in a 37°C water bath before testing the activity.

### Kinase Assays

CheA kinase activity was measured using an enzyme-coupled assay that couples ATP consumption to NADH oxidation, which is monitored by the decrease of NADH absorbance at 340 nm for 90 seconds (8). 2 µL of the complex was mixed with 198 µL of a CheY mixture that contained 55 µM CheY, 20 units of enzymes (lactate dehydrogenase (LDH) and pyruvate kinase (PK)) (Sigma-Aldrich), 4 mM ATP, 2.2 mM phosphoenolpyruvate (PEP), and 250 µM NADH in kinase buffer at pH 7.4. The adjusted kinase activity was calculated by subtracting the slope of the absorbance change at 340 nm measured for the CheY-only background control from the slope measured for the complex. Specific kinase activity per bound CheA (s^-1^) was calculated as follows: [(adjusted slope as abs/sec) x (M NADH/6220 abs)] / [(M CheA in complex) x (2 µL/200 µL)]. CheA concentration in the complex was determined by the sedimentation assay described below.

### Protein sedimentation assays

Sedimentation assays were used to determine amount of protein bound to vesicles to form functional complexes. 40 µL of functional complexes was spun in tabletop ultracentrifuge (Beckman TLX, TLA 120.2 rotor, 60,000 rpm) at 25°C for 30 min. Prior to sedimentation, a gel sample was prepared corresponding to the total amount of CF, CheA, and CheW in the sample. After sedimentation, the supernatant which contained unbound CF, CheA and CheW was removed and placed in a clean Eppendorf tube and used to prepare a supernatant gel sample. The pellet was resuspended to the original volume using kinase buffer, vortexed, and used to prepare a pellet gel sample. Total, supernatant and pellet gel samples were run on 12.5 or 15 % acrylamide SDS-PAGE along with gel samples that contained known concentrations of CF, CheA and CheW for calibration. Gels were then imaged by densitometry with a Gel Doc EZ Imager (Bio-Rad) and were analyzed with ImageJ software (42). Band intensities of proteins in three lanes (dilution series of the known concentration gel sample) were used to obtain linear fit parameters for CheA, CF, and CheW to correct for the gel background and staining differences, and calculate the protein concentrations. The stoichiometry of the proteins in the array was calculated based on (1) the ratio of the concentrations calculated from the total minus super ([*Protein*]_Total_ x [*I*_Total_-*I*_Supernatant_]/I_total_), and (2) the ratio of protein concentrations in the pellet fraction. The resulting stoichiometries from both methods were similar and were averaged to determine the reported stoichiometry.

### HDX Mass Spectrometry Measurements

#### HDX sample preparation

Vesicle-mediated complexes (CF4Q, CF4Q.A411V or CF4E with CheA and CheW) were assembled and incubated overnight, followed by kinase activity and protein binding measurements to be sure complexes were active and to monitor the amount of CheA and CheW bound to CF throughout the time-course of exchange. Before initiating exchange, a 2 mL G10 Sephadex Zeba desalting column (Pierce Biotechnology) was pre-equilibrated with deuterated 1x kinase buffer (50 mM K_x_H_x_PO_4_, 50 mM KCl, and 5 mM MgCl_2_ at pD 7.5 corresponding to a pH reading of 7.1) at 25°C as follows: spin column at 2000 rpm for 2 min in a tabletop centrifuge (Beckman Coulter Allegra R Tabletop Centrifuge), discard the flow-through, add 1 mL deuterated buffer to the top of the column, spin again and repeat until the column has been exchanged with a total of 4 ml of deuterated buffer. Exchange was initiated by adding 1 mL of complexes (assembled on vesicles with 30 µM CF, 12 µM CheA, 24 µM CheW) to the pre-equilibrated column followed by centrifugation at 2000 rpm for 2 min at 25°C. The eluted solution of complexes was incubated in a 25°C water bath. For each time point of exchange, 22.5 µL was transferred into a pre-chilled Eppendorf tube containing 22.5 µL quench buffer (1% formic acid, 20% w/v glycerol, 1M GuHCl, pH 1.6) in a 0°C ice-water bath. Each sample was immediately vortexed, flash-frozen in liquid nitrogen, and stored at -80°C. Undeuterated samples were prepared in the same manner, without the buffer exchange step on the column (22.5 µL of the initial assembly added to 22.5 µL of pre-chilled quench buffer and then flash-frozen).

The reversible heat denaturation of CF was used for preparation of a fully-exchanged sample (11), which was subjected to the mass spectrometry protocol to measure the back exchange for each peptide for our protocol. The fully-exchanged sample was prepared by applying 1 mL of a 30 µM CF4Q solution to a desalting column for buffer exchange into D_2_O (as described above). The exchanged CF4Q was then placed in an 80°C water bath for denaturation for 1 hour, followed by 30 min in a 0°C ice-water bath. The sample was prepared as described above (22.5 µL added to 22.5 µL of pre-chilled quench buffer and then flash-frozen).

#### HPLC column preparation and maintenance

For the MS analysis, a high performance liquid chromatography (HPLC) column was used for desalting and peptide separation. Before each set of mass spectrometry experiments, the HPLC column (2.1 mm x 5 cm C18 reverse phase column from Higgins Analytical) was connected to an LC pump (Agilent 1100G1312A) and cleaned by flowing 200 µL/min of Buffer B (0.1% formic acid in acetonitrile) overnight. At the end of each mass spectrometry experiment, the column was stored with 100% Buffer B. Every two months, the column was subjected to deep cleaning by flushing with 20 column volumes of water, 20 column volumes of acetonitrile, 5 column volumes of isopropanol, 20 column volumes of heptane, 5 column volumes of isopropanol, and then 20 column volumes of acetonitrile at 200 µL/min.

### HDX-MS data acquisition

In order to minimize back-exchange, the HPLC separation was performed at low temperature. The column, connecting tubing and valves were all maintained near 0 °C using an ice box filled with ice water and the solvent was pre-chilled. Valves and lines were thoroughly cleaned prior to use.

Immediately prior to MS analysis, the HPLC column was equilibrated with 95 % Buffer A (0.1% formic acid in water). Samples of frozen exchanged complexes were thawed for 1 min in a 0°C ice water bath, pepsin (Sigma Aldrich) was added in a 1:1 protein:enzyme molar ratio for 1 min of digestion at 0°C, and then 45 µL was immediately injected via the injection valve on to the chilled column. Peptides were eluted at 100 µL/min directly into the Waters Synapt G2Si with the following gradient: 5% Buffer B (0.1% formic acid in acetonitrile) for 3 min for desalting; from 5-50% B over 12 min to separate peptides, hold at 50% B for 3 min to elute all remaining peptides, 95% B for 7 min to clean the column, 5% B for 15 min to re-equilibrate the column ready for the next injection. Only the portion 3-18 min was injected into the mass spectrometer; the initial 3 min (containing salts) and the 18-40 min cleaning (containing lipids) and re-equilibration were diverted to waste to minimize ion source contamination.

At least one undeuterated sample was run on each day of MS experiments with collision-induced dissociation in MS^E^ mode to confirm peptide identification. One fully-exchanged control sample was run each day of experiments to measure back exchange. Water blanks were run between each sample to ensure no carryover from the previous sample. All exchanged samples were run in random order.

The mass spectrometer was calibrated each day in positive ion mode using sodium formate. All samples were acquired in positive ion mode with ion mobility separation in the mass range 100-2000 m/z. A 100 fmol/µL solution of leucine enkephalin ([M+H]^+^ 556.2771 m/z) was continuously infused at 2 µL/min for lock mass correction.

### HDX data analysis

Peptides were identified by ProteinLynx Global Server (PLGS) version 3.0.1 software from Waters. MS^E^ mass spectral data acquired from undeuterated samples were searched against a database containing the sequences of proteins in the complexes. Process parameters were set to using 556.2771 Da/e as lock mass for charge 1, lock mass window as 0.25 Da with non-specific primary digest reagent. Peptide results generated by PLGS were then imported to DynamX version 3.0 software for further analysis.

Peptide results were imported into DynamX with the following parameters: minimum signal intensity of 5000, minimum products per amino acid of 0.3, maximum sequence length of 25 amino acids, retention time RSD of 2%. Peptides were manually checked to be sure they each contained a minimum of 3 consecutive product ions. Data were manually curated after automatic analysis by checking that each isotope peak in the isotopic distribution for a peptide had the same retention time and ion mobility, that different charge states of the same peptide had the same retention time, and that the same peptide in different samples exhibited similar retention times and ion mobility. The deuterium incorporation of each peptide was determined by subtracting the centroid mass of the undeuterated sample from that of the deuterated sample at each time point. The threshold for a significant difference between HDX uptake for two states was calculated as follows (43): a simple average of the standard deviations for the calculated means of all the data for two samples was divided by the square root of the number of data sets (2) to yield the standard error of the mean, which was multiplied by the Student’s t-table value for a 95% confidence interval and two degrees of freedom. The resulting threshold for a significant difference was ± 0.8 or ± 0.9 for the comparisons shown in Figures 4-5.

EX1 analysis was initially performed using HX-Express2 (19) to deconvolute bimodal patterns into two distributions. Chosen parameters include a distribution width of 20% of BPI (base peak intensity), isotopic peak detection with peak tolerances of 5% of BPI, and 3 data points, and 4 channel Savitsky-Golay smoothing. Spectra exhibiting a bimodal distribution were analyzed by fitting to a double binomial by providing the number of amides of the peptide and specifying a fitting asymmetry of 1 to obtain the deconvolution plots and the probability and relative intensity of both low and high mass distributions.

## Acknowledgments

This research was supported by National Institutes of Health Grant R01-GM085288 and R01-GM120195. Mass spectrometry data were obtained at the University of Massachusetts Mass Spectrometry Facility, with support from the Institute for Applied Life Sciences. We thank Brian Crane for providing coordinates for the structural model.

## ^1^Abbreviations

CF: cytoplasmic fragment of *E. coli* Asp receptor;
DOPC: 1,2-dioleoyl-sn-glycero-3-phosphocholine;
DGS-NTA: 1,2-dioleoyl-sn-glycero-3-[[N(5-amino-1-carboxypentyl)iminodiacetic acid]succinyl];
ESI: electrospray ionization;
HAMP domain: present in Histidine kinases, Adenylate cyclases, Methyl accepting proteins and Phosphatases;
HDX-MS: hydrogen deuterium exchange mass spectrometry;
IPTG: isopropyl β-D-1-thiogalactopyranoside;
LB: Luria-Bertani;
MH1: methylation helix 1;
MH2: methylation helix 2;
PMSF: phenylmethane sulfonyl fluoride;
PF: protection factor;
TEV: tobacco etch virus.

