## Supplemental Information for "Hydrogen exchange of chemoreceptors in functional complexes suggests protein stabilization mediates long-range allosteric coupling"

**Supporting Information**

### Hydrogen exchange properties of bacterial chemotaxis receptors in functional complexes suggest an order/disorder transition mechanism of signaling

Xuni Li, Stephen J. Eyles, and Lynmarie K. Thompson

**Table S1.** Kinase activity and protein incorporation into functional complexes of CF4Q, CF4Q.A411V, and CF4E, assembled on vesicles with CheA and CheW.

| Time | Specific Activity[a] ($s^{-1}$) | CF[b] (μM) | CheA[b] (μM) | CheW[b] (μM) | CF:CheA:CheW[c] |
|---|---|---|---|---|---|
| | | | **CF4Q** | | |
| **0min** | 23.3 ±5 | 24.3 ±2 | 3.7 ±0.1 | 10.8 ±0.1 | 6: 1: 2.7 |
| **960min (16 hr)** | 26 ±3 | 27 ±1.3 | 4.75 ±0.4 | 13.2 ±3.7 | 6: 1.2: 2.8 |
| | | | **CF4Q.A411V** | | |
| **0min** | 0.04 ±0.14 | 24.4 ±0.5 | 3.75 ±0.2 | 11.6 ± 1.8 | 6:1:2.7 |
| **960min (16 hr)** | 0.01 ±0.11 | 28.4 ±5.2 | 4.85 ±1.2 | 13.5 ±0.7 | 6:1.3:2.9 |
| | | | **CF4E** | | |
| **0min** | 17.9 ±4 | 26.3 ±7.3 | 3.4 ±0.1 | 9.6 ±0.4 | 6:0.9:2.3 |
| **960min (16 hr)** | 11.2 ±3.8 | 29.3 ±3 | 5.3 ±1.6 | 12.6 ±1 | 6:1:2.6 |

[a]Activities are average of four replicates (measured on two samples on two different days); error bars indicate ± one standard deviation.
[b]Protein concentrations in the sedimented complexes (after respuspension to original volume), based on SDS-PAGE band intensities.
[c]Stoichiometry of proteins in the sedimented complex was calculated by setting the CF concentration (24-29 μM) to 6 to calculate the ratio of CF to CheA and CheW.

**Table S2**. Best fit parameters[a] for HDX uptake (curves shown in Figure S5) for representative peptides throughout CF.

| | | | **fast phase** | | | **slow phase** | | | |
|---|---|---|---|---|---|---|---|---|---|
| **peptide**[b] | **state** | $k_{int}$[c] ($min^{-1}$) | $f_1$ | $k_1$ ($min^{-1}$) | **log(PF)**[c] | $f_2$ | $k_2$ ($min^{-1}$) | **log(PF)**[c] | $k_{avge}$[d] ($min^{-1}$) |
| 260-273 | 4Q | 500.49 | | | | | | | |
| | 4E | 500.49 | 1.00 | ≥ 2.00 | 2.4 | 0.00 | | | 2.00 |
| | 4QA411V | 500.49 | | | | | | | |
| 308-326 | 4Q | 792.97 | 0.74 | ≥ 0.50 | 3.2 | 0.26 | 0.041 | 4.3 | 0.38 |
| | 4E | 769.30 | **0.85** | ≥ 0.84 | 3.0 | 0.15 | 0.089 | 3.9 | 0.73 |
| | 4QA411V | 792.97 | **0.85** | ≥ 0.70 | 3.1 | 0.15 | 0.044 | 4.3 | 0.60 |
| 328-351 | 4Q | 680.43 | 0.65 | 0.42 | 3.2 | 0.35 | 0.021 | 4.5 | 0.28 |
| | 4E | 680.43 | **0.78** | ≥ 0.71 | 3.0 | 0.22 | 0.050 | 4.1 | 0.56 |
| | 4QA411V | 680.43 | 0.58 | 0.38 | 3.3 | 0.42 | 0.020 | 4.5 | 0.23 |
| 362-368 | 4Q | 305.85 | 0.63 | 0.37 | 2.9 | 0.37 | 0.036 | 3.9 | 0.25 |
| | 4E | 305.85 | **0.72** | ≥ 0.77 | 2.6 | 0.28 | 0.041 | 3.9 | 0.57 |
| | 4QA411V | 305.85 | **0.79** | 0.41 | 2.9 | 0.21 | 0.006 | 4.7 | 0.32 |
| 369-376 | 4Q | 684.94 | 0.30 | 0.19 | 3.6 | 0.70 | 0.006 | 5.1 | 0.06 |
| | 4E | 684.94 | **0.48** | 0.43 | 3.2 | 0.52 | 0.006 | 5.0 | 0.21 |
| | 4QA411V | 684.94 | 0.25 | 0.22 | 3.5 | 0.75 | 0.009 | 4.9 | 0.06 |
| 377-383 | 4Q | 290.77 | 0.00 | | | 1.00 | 0.006 | 4.7 | 0.01 |
| | 4E | 290.77 | **0.34** | ≥ 0.90 | 2.5 | 0.66 | 0.004 | 4.9 | 0.31 |
| | 4QA411V | 290.77 | 0.00 | | | 1.00 | 0.007 | 4.6 | 0.01 |
| 384-394 | 4Q | 293.99 | 0.92 | ≥ 0.69 | 2.6 | 0.08 | 0.040 | 3.9 | 0.64 |
| | 4E | 293.99 | 0.90 | ≥ 1.07 | 2.4 | 0.10 | 0.016 | 4.3 | 0.97 |
| | 4QA411V | 293.99 | 0.87 | ≥ 1.04 | 2.4 | 0.13 | 0.025 | 4.1 | 0.91 |
| 395-417 | 4Q | 666.35 | 0.36 | 0.26 | 3.4 | 0.64 | 0.006 | 5.0 | 0.10 |
| | 4E | 666.35 | | | | | | | |
| | 4QA411V | 630.22 | 0.36 | 0.23 | 3.4 | 0.64 | 0.005 | 5.1 | 0.08 |
| 418-424 | 4Q | 370.59 | 0.65 | ≥ 0.38 | 3.0 | 0.35 | 0.023 | 4.2 | 0.25 |
| | 4E | 370.59 | 0.59 | ≥ 1.10 | 2.5 | 0.41 | 0.103 | 3.6 | 0.69 |
| | 4QA411V | 370.59 | 0.61 | ≥ 0.52 | 2.9 | 0.39 | 0.045 | 3.9 | 0.34 |
| 422-432 | 4Q | 312.90 | 0.65 | ≥ 0.67 | 2.7 | 0.35 | 0.149 | 3.3 | 0.49 |
| | 4E | 312.90 | **0.91** | ≥ 1.05 | 2.5 | 0.09 | 0.116 | 3.4 | 0.97 |
| | 4QA411V | 312.90 | 0.65 | ≥ 0.67 | 2.7 | 0.35 | 0.150 | 3.3 | 0.49 |
| 433-448 | 4Q | 879.21 | 0.76 | 0.55 | 3.2 | 0.24 | 0.016 | 4.7 | 0.42 |
| | 4E | 879.21 | **0.81** | ≥ 0.68 | 3.1 | 0.19 | 0.021 | 4.6 | 0.55 |
| | 4QA411V | 879.21 | 0.75 | 0.40 | 3.3 | 0.25 | 0.012 | 4.9 | 0.30 |
| 447-456 | 4Q | 449.27 | 0.52 | 0.36 | 3.1 | 0.48 | 0.014 | 4.5 | 0.19 |
| | 4E | 449.27 | **0.65** | 0.51 | 2.9 | 0.35 | 0.013 | 4.5 | 0.33 |

| | | | | | | | | | |
|---|---|---|---|---|---|---|---|---|---|
| | 4QA411V | 449.27 | 0.52 | 0.45 | 3.0 | 0.48 | 0.021 | 4.3 | 0.24 |
| 455-470 | 4Q | 703.64 | 1.00 | 0.77 | 3.0 | 0.00 | | | 0.77 |
| | 4E | 703.64 | 1.00 | ≥ 1.15 | 2.8 | 0.00 | | | 1.15 |
| | 4QA411V | 703.64 | 1.00 | 0.95 | 2.9 | 0.00 | | | 0.95 |
| 471-478 | 4Q | 355.03 | 0.59 | 0.53 | 2.8 | 0.41 | 0.013 | 4.4 | 0.32 |
| | 4E | 355.03 | **0.76** | ≥ 0.87 | 2.6 | 0.24 | 0.009 | 4.6 | 0.66 |
| | 4QA411V | 355.03 | **0.72** | 0.51 | 2.8 | 0.28 | 0.009 | 4.6 | 0.36 |
| 478-488 | 4Q | 316.13 | 0.76 | 0.53 | 2.8 | 0.24 | 0.029 | 4.0 | 0.41 |
| | 4E | 316.13 | 0.77 | ≥ 0.93 | 2.5 | 0.23 | 0.014 | 4.4 | 0.72 |
| | 4QA411V | 316.13 | **0.81** | 0.56 | 2.8 | 0.19 | 0.009 | 4.5 | 0.46 |
| 489-499 | 4Q | 282.17 | 0.72 | 0.55 | 2.7 | 0.28 | 0.016 | 4.2 | 0.40 |
| | 4E | 263.27 | **0.78** | 0.64 | 2.6 | 0.22 | 0.011 | 4.4 | 0.50 |
| | 4QA411V | 282.17 | **0.79** | 0.62 | 2.7 | 0.21 | 0.012 | 4.4 | 0.49 |
| 500-512 | 4Q | 672.50 | 0.70 | 0.52 | 3.1 | 0.30 | 0.023 | 4.5 | 0.37 |
| | 4E | 672.50 | **0.77** | ≥ 0.98 | 2.8 | 0.23 | 0.021 | 4.5 | 0.76 |
| | 4QA411V | 672.50 | **0.77** | 0.59 | 3.1 | 0.23 | 0.013 | 4.7 | 0.45 |
| 526-532 | 4Q | 315.23 | | | | | | | |
| | 4E | 315.23 | 1.00 | 0.76 | 2.6 | 0.00 | | | 0.76 |
| | 4QA411V | 315.23 | 1.00 | 0.46 | 2.8 | 0.00 | | | 0.46 |
| [e]Average $f_1$ | 4Q | | 0.7 | | | | | | |
| | 4E | | 0.8 | | | | | | |
| | 4QA411V | | 0.7 | | | | | | |
| | | | | PF range: | ≤2.4-3.6 | | PF range: | 3.3-5.1 | |
| | | | | Avge: | 2.9 | | Avge: | 4.4 | |

[a]Data were fit to the equation $y = p_1 + p_2 - \left(p_1 e^{-k_1 t} + p_2 e^{-k_2 t}\right)$ for biexponential uptake, to obtain the population and rate constant of fast ($p_1$, $k_1$) and slow uptake ($p_2$, $k_2$). Fractions of each phase are reported: $f_1 = \frac{p_1}{p_1+p_2}$ and $f_2 = \frac{p_2}{p_1+p_2}$. A few peptides were well fit with a monoexponential curve. All curves plotted in Figure S5.

[b]Protein regions for each peptide are color coded. Red represents regions that exhibit very rapid HDX, not resolved by our experiment: methylation helix 1 (263-318), cytoplasmic tip (384-394), and cytoplasmic tail (516-551). Blue represents regions that exhibit slow HDX: the protein interaction region, excluding the cytoplasmic tip (361-383 and 395-417). Purple represents regions that exhibit intermediate HDX properties: flexible bundle (319-360 and 418-459) and methylation helix 2 (460-515). Note that peptide 308-326 (represented in purple) spans both methylation helix 1 and flexible bundle regions.

[c]The intrinsic exchange rate for each peptide, $k_{int}$, was calculated using the SPHERE program (https://landing.foxchase.org/research/labs/roder/sphere/) for pDcorr=7.5 and 25˚C. Protection factors (PF) for the fast and slow phases are calculated as PF $=k_{int}/k_1$ and PF $=k_{int}/k_2$, respectively. For uptake data that included a very rapid phase that was not well defined by the data (first time point at 3 min), $k_1$ is a lower limit and the protection factor, listed in red, is an upper limit. For these cases, a larger $k_1$ fit equally well, as shown by the dotted/dashed curves in Figure S5. The last rows give the range and average log(PF) for the fast and slow phase.

[d]Average rate $k_{avg} = f_1 * k_1 + f_2 * k_2$.

[e]Average fraction fast phase for peptides where all 3 states exhibit a fast phase. Individual values of $f_1$ for 4E and 4Q.A411V that differ from 4Q are highlighted in bold

**Table S3.** Receptor residues interacting with CheA and CheW based on this and other studies.

| *Method: proteins present*[a] | *Receptor Residues Involved in Interaction*[b] | *Ref* |
|---|---|---|
| **Receptor/CheA interface** | | |
| NMR: $TM0014_{90-206}$[c] , CheAΔ354[d] | 375, 376, 378, 382[h] | (1) |
| Disulfide crosslinking: **Tsr, CheA, CheW** | 381, **385**, 396, 399[i] | (2) |
| X-ray (3UR1): Tm14s[e] , CheAΔ354[d] , CheW<br>ECT: **native E coli receptors, CheA, CheW** | 371, 375, 378, 379, 382, **386**, 400, 406, 410[j] | (3) |
| ECT & MD (3JA6): **Tar-CF, CheA, CheW**[f] | **369**, 375, 378, 379, **392**[j] | (4) |
| **Receptor/CheW interface** | | |
| NMR: $TM0014_{90-206}$[c] , CheW | 375, 376, 378, 382[h] | (1) |
| NMR: $TM0014_{90-206}$[c] , CheW | 372, 377, 379, 380–383, **385**, **386**, 396[k] | (5) |
| Disulfide crosslinking: **Tsr, CheA, CheW** | **386**, 396[l] | (6) |
| X-ray (3UR1): Tm14s[e] , CheAΔ354[d] , CheW<br>ECT: **native E coli receptors, CheA, CheW** | 378, 379, 381, 383[m] | (3) |
| ECT & MD (3JA6): **Tar-CF, CheA, CheW**[f] | 375, 378, 379, 382, 383, **385**, **386**, **392**, 402[m] | (4) |
| **Receptor/CheA or CheW interface** | | |
| HDX-MS: **Tar-CF, CheA, CheW**[g] | Residues within 372-383, 395-417[n] | this study |

[a]Only the protein complexes highlighted in bold green are functional
[b]Asp receptor (*E. coli* Tar) residue numbers are obtained as follows: TM0014 number + 240 = Tar number; Tm14s number - 2002 = Tar number; Tsr number - 2 = Tar number. Bold red indicates contacts reported in previous studies that are not within the regions with incomplete HDX at 16 hr.
[c]$TM0014_{90-206}$ is a truncated version of TM0014, a single chain soluble chemoreceptor which lacks the transmembrane region.
[d]CheAΔ354 is a monomeric construct containing only P4 and P5
[e]Residues 107-191 of the *T maritima* receptor Tm14s, corresponding to the protein interaction domain.
[f]Functional complexes assembled on lipid monolayers.
[g]Functional complexes assembled on lipid vesicles.
[h]Chemical shift perturbation of receptor $^{13}$C-methyl ILV sidechains upon binding
[i]In vitro crosslinking of Cys mutants
[j]Residues of Tar that are within 3Å of CheA in Crane model or 3JA6.
[k]Chemical shift perturbation of receptor $^{15}$N-amide backbone upon binding
[l]In vivo crosslinking of Cys mutants
[m]Residues of Tar that are within 3Å of CheW in Crane model or 3JA6.
[n]Incomplete HDX at 16 hr.

#### A. CF4Q Peptide Coverage

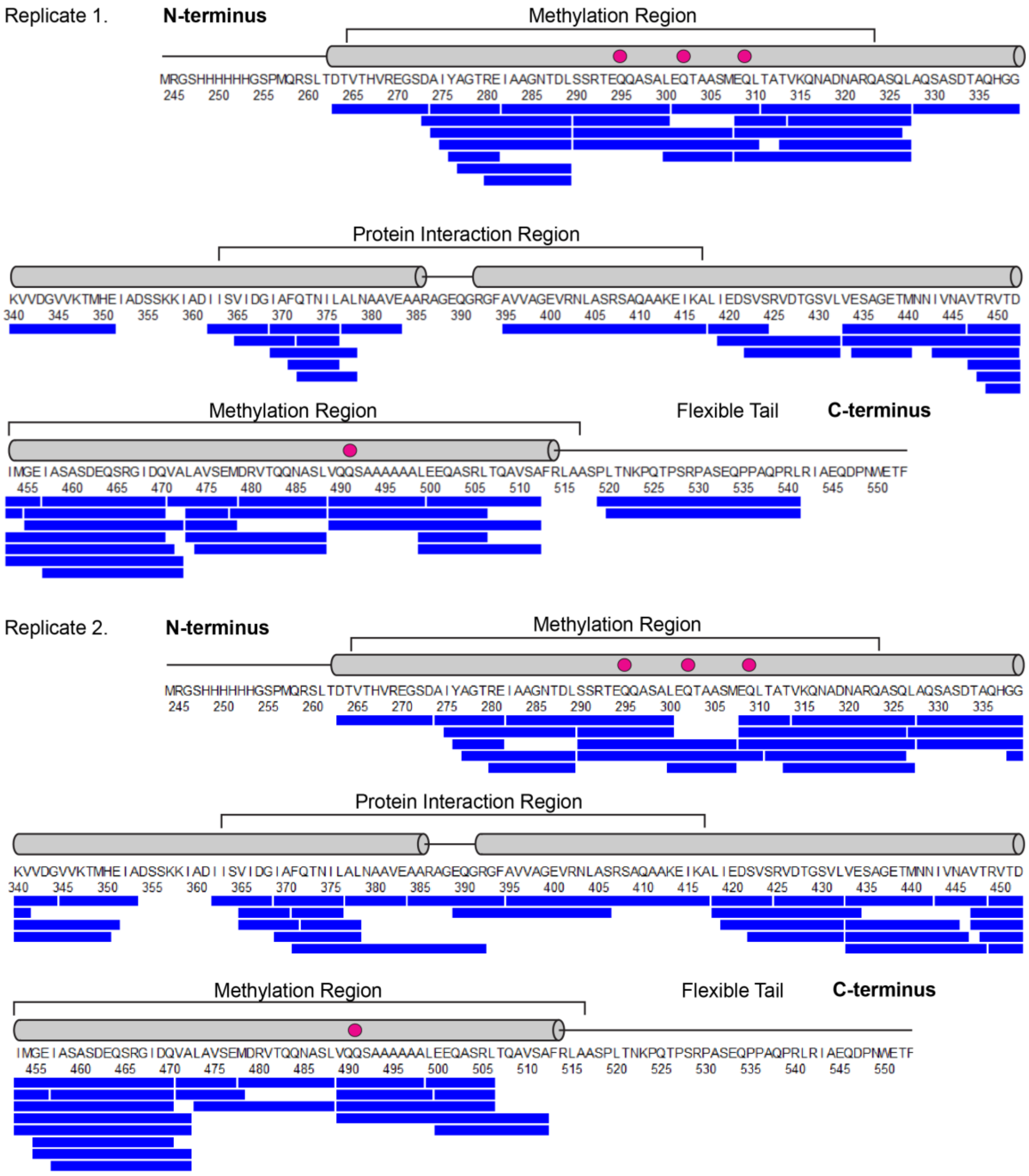

#### B. CF4Q.A411V Peptide Coverage

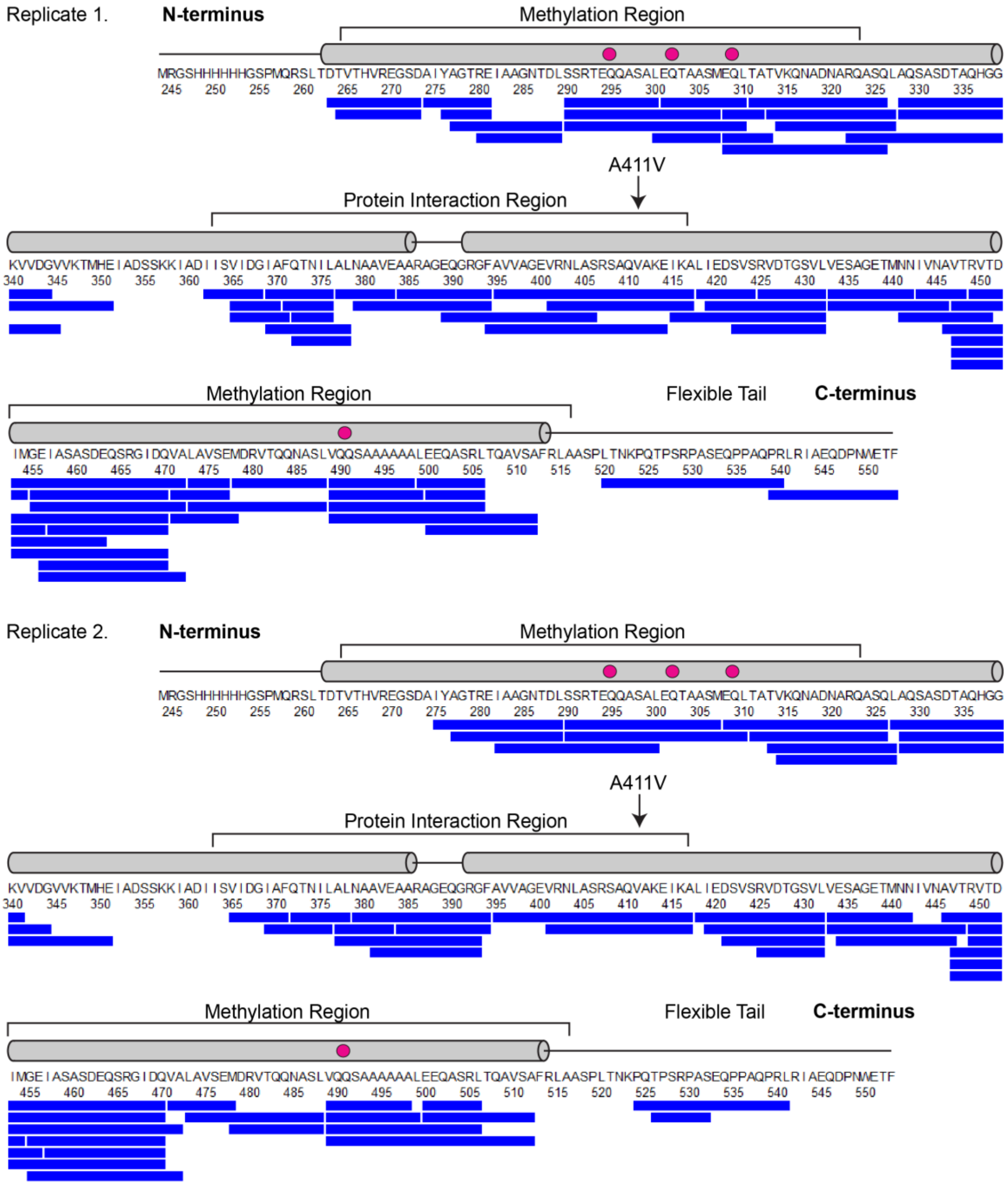

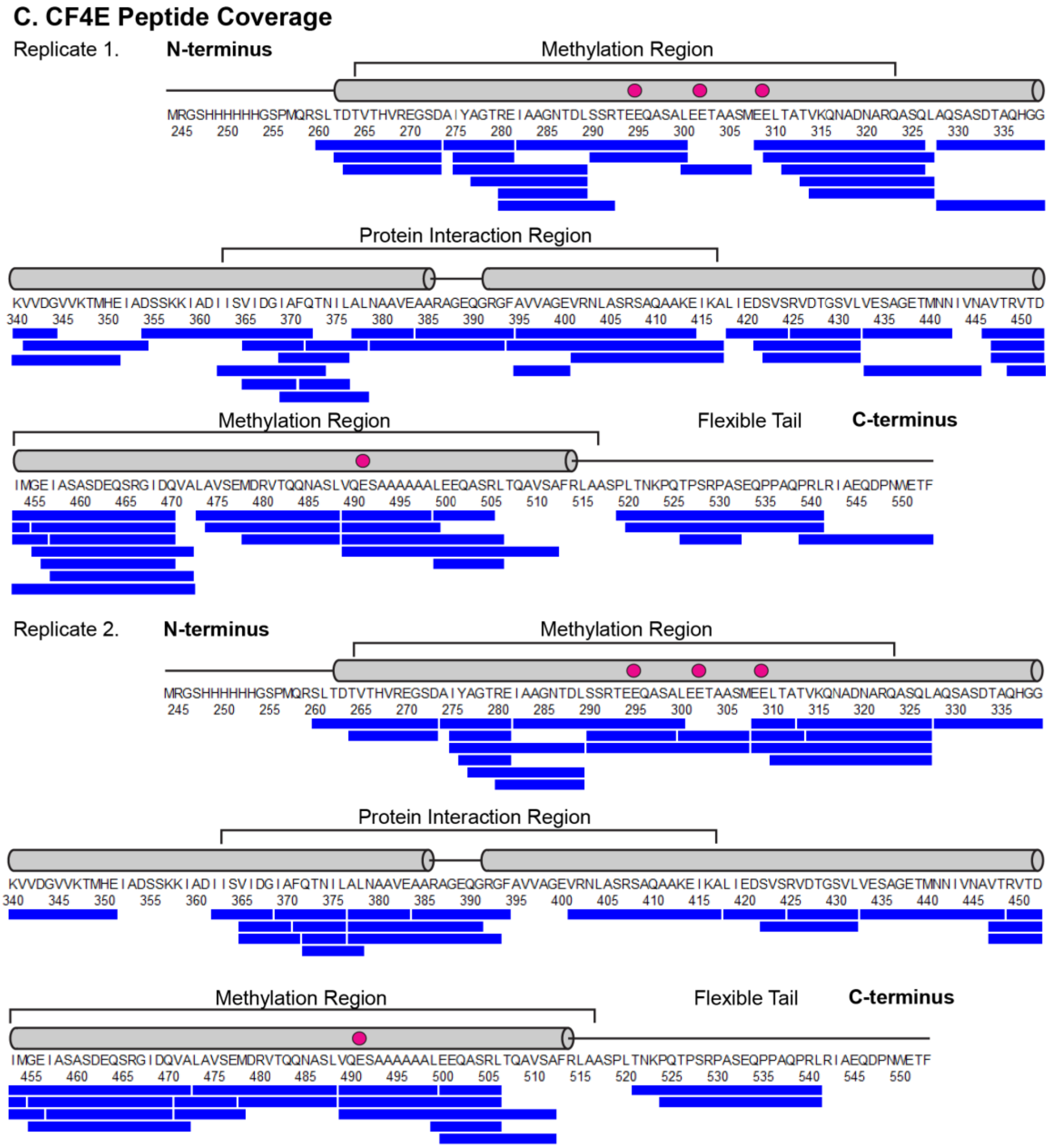


**Figure S1.** Peptide coverage maps for CF in functional complexes with CheA and CheW. Peptide numbering corresponds to the sequence of the intact *E. coli* Asp receptor. Blue bars below the sequence are CF peptides from each of two replicates for (A) CF4Q, (B) CF4Q.A411V and (C) CF4E. CF secondary structure elements and regions are labelled above the sequence, and methylation sites are shown as magenta circles. Average peptide coverage and amino acid redundancy are 80% and 3.4 for CF4Q, 83% and 3.1 for CF4Q.A411V, and 88% and 2.6 for CF4E.

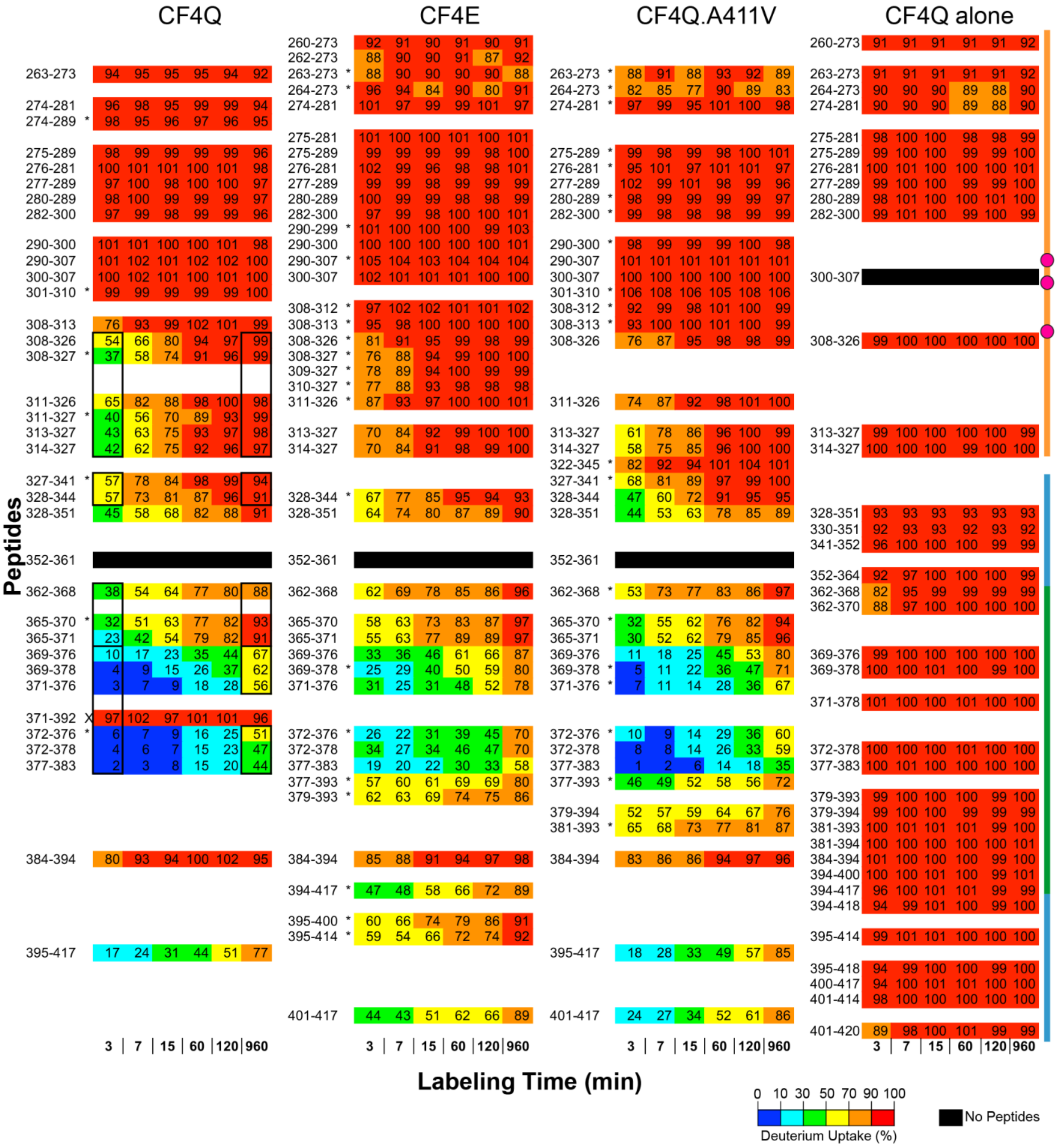
CF4Q
CF4E
CF4Q.A411V
CF4Q alone
Peptides
Labeling Time (min)
3 | 7 | 15 | 60 | 120 | 960
0 10 30 50 70 90 100
Deuterium Uptake (%)
No Peptides

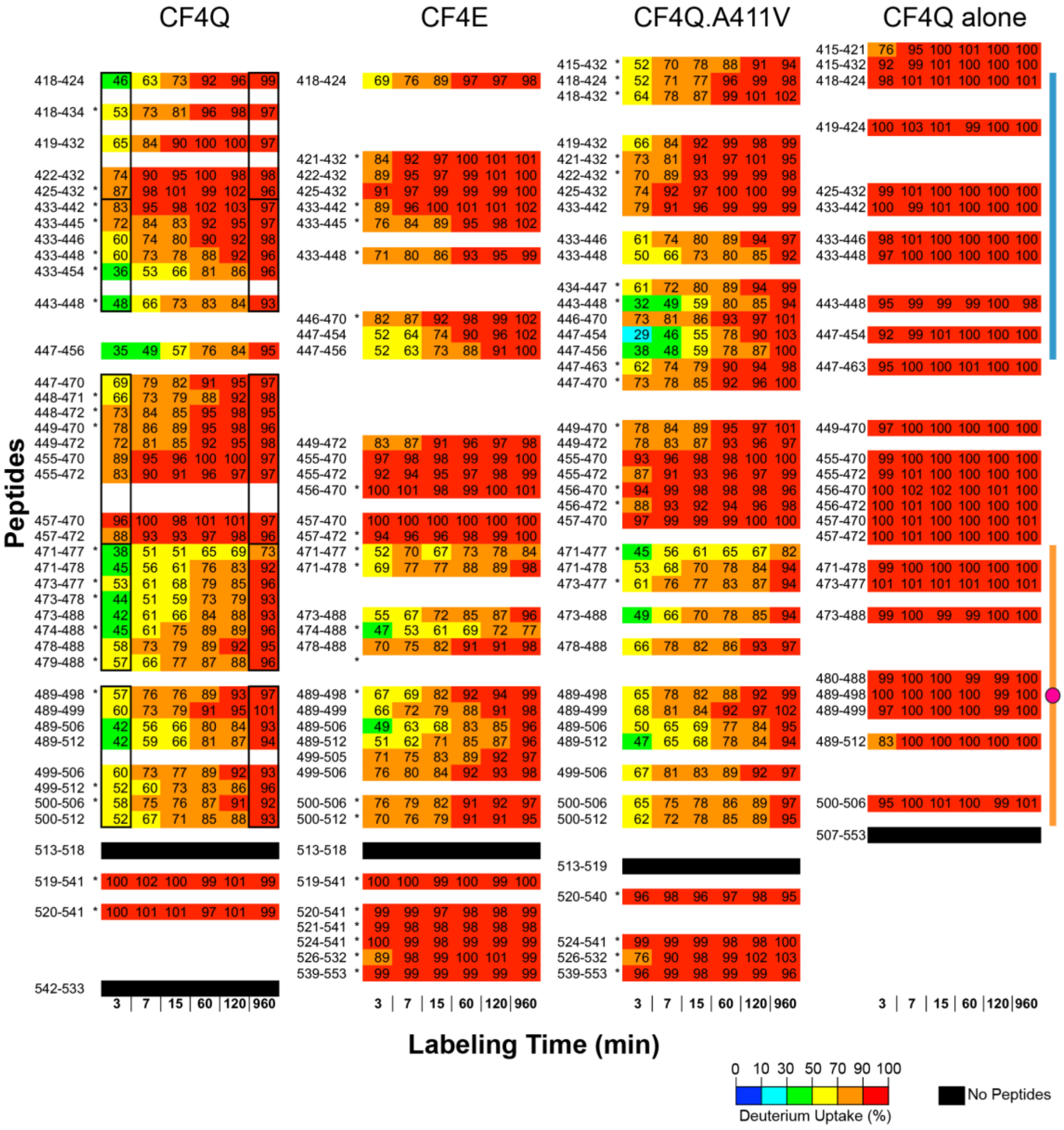


**Figure S2**. Hydrogen exchange for the complete set of CF peptides from CF4Q, CF4E, and CF4Q.A411V in functional complexes with CheA and CheW, and from CF4Q alone. A subset of the CF4Q data are shown in Figure 2. Percent deuterium uptake is calculated based on a centroid analysis, as the uptake of the peptide at each time point divided by the uptake of the fully exchanged sample, and is represented with both numbers and rainbow colors from blue to red for low to high uptake. Regions where no peptides were observed are shown in black. Peptides with asterisks were found in a single data set; others were found in both replicates. Finally, X marks an outlier peptide (371-392) that we neglected in the analysis because a single replicate for a single state was observed and its rapid exchange was inconsistent with the slow exchange of six overlapping peptides. Uptake for overlapping peptides outlined by black boxes was averaged to choose colors represented on Figure 2 for 3 min and 16 hour HDX. Note that for the sample of CF4Q alone, nearly all peptides exhibit ≥ 90% exchange (red) within 3 min. The few time points that show 70-90% exchange (orange) all have at least one overlapping peptide with ≥ 90% exchange (or 88% exchange for residues 365-368). Thus, we conclude that the entire CF alone sample exhibits very rapid exchange (≥ 90% within 3 min). Interestingly, one region (328–351) outside of the protein interaction region exhibits incomplete exchange at a significant level in both functional complexes and CF4Q alone (1.6–1.9 Da unexchanged at 16 hr in peptide 328-351 of CF4Q, CF4E, and CF4Q.A411V in complexes, and 1.4 Da unexchanged at 16 hr for peptides 328–351 and 330–351 of CF4Q alone). As explained in the Methods section, differences greater than 0.9 Da are judged to be significant. We conclude that incomplete exchange in the 328–351 region is an intrinsic property of the CF protein itself, and all other sites of incomplete exchange in CF complexes localize to the protein interaction region. Vertical lines on the right side of the figure represent regions of the CF, in colors consistent with Figure 3: orange lines indicate the methylation region, with methylation sites shown as magenta circles, blue lines indicate the flexible bundle region, and the green line indicates the protein interaction region.

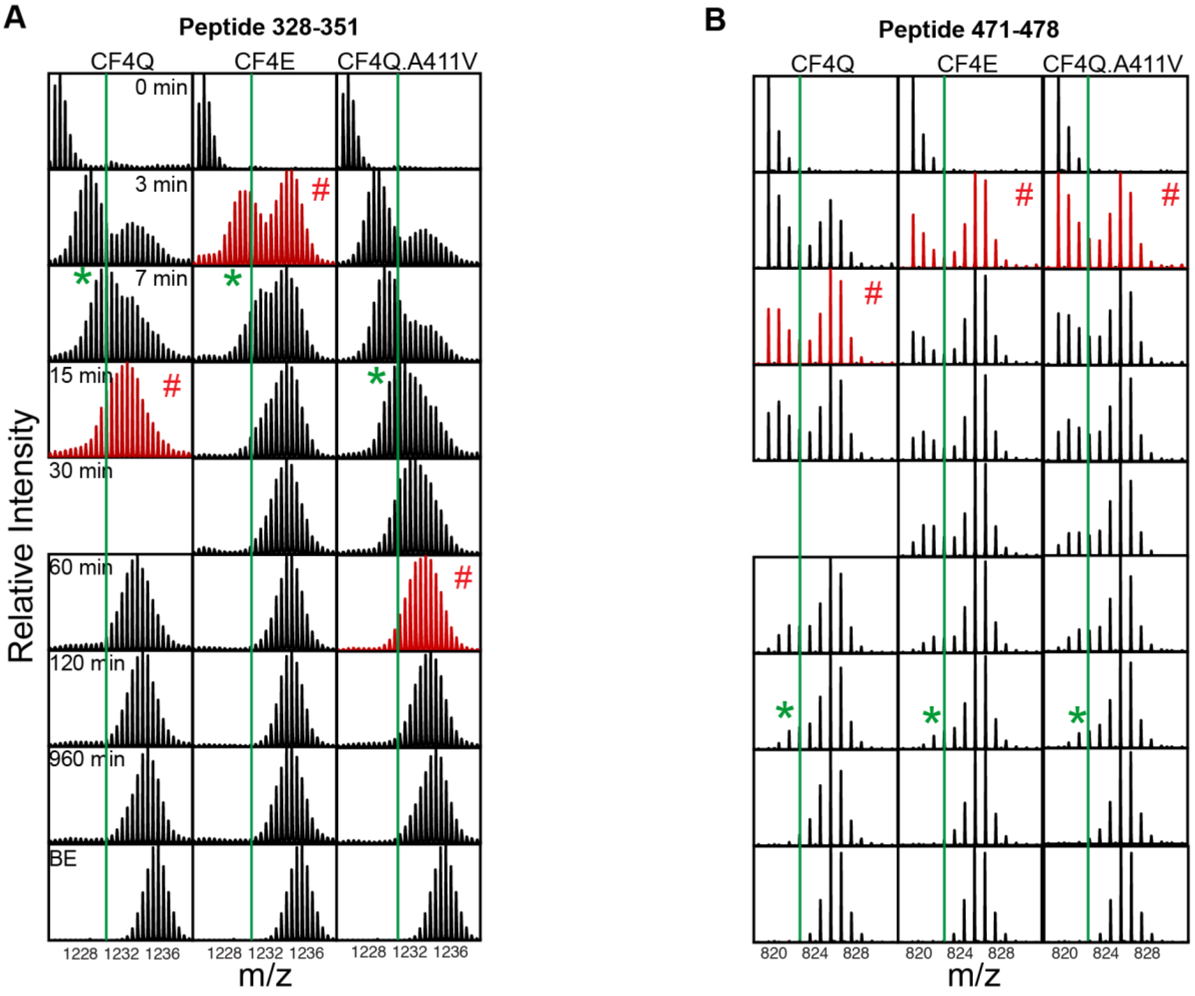


Relative intensity of high m/z distribution, based on HX-Express deconvolutions. Bold red indicates $t_{1/2}$ of EX1 process has been reached.

| | Peptide 328-351 | | | Peptide 471-478 | | |
|---|---|---|---|---|---|---|
| **Time (min)** | **CF4Q** | **CF4E** | **CF4Q.A411V** | **CF4Q** | **CF4E** | **CF4Q.A411V** |
| **3** | 0.20 | **0.55** | 0.32 | 0.46 | **0.72** | **0.56** |
| **7** | 0.29 | 0.66 | 0.29 | **0.52** | 0.78 | 0.64 |
| **15** | **0.34** | 0.46 | 0.34 | 0.6 | 0.76 | 0.69 |
| **30** | NA | 0.81 | 0.42 | NA | 0.74 | 0.78 |
| **60** | NA | 0.84 | **0.54** | 0.69 | 0.84 | 0.84 |
| **120** | NA | 0.96 | 0.75 | 0.76 | 0.84 | NA |

Uptake of low m/z distribution, based on HX-Express deconvolutions. Bold green indicates $t_{1/2}$ of EX2 process has been reached.

| | Peptide 328-351 | | | Peptide 471-478 | | |
|---|---|---|---|---|---|---|
| **Time (min)** | **CF4Q** | **CF4E** | **CF4Q.A411V** | **CF4Q** | **CF4E** | **CF4Q.A411V** |
| **3** | 7.31 | 8.48 | 5.53 | 0.31 | 0.3 | 0.49 |
| **7** | **10.31** | **10.92** | 7.29 | 0.47 | 0.48 | 0.76 |

| | | | | | | |
|---|---|---|---|---|---|---|
| **15** | 11.48 | 13.53 | **9.5** | 0.78 | 0.83 | 1.06 |
| **30** | NA | 13.31 | NA | NA | 1.34 | NA |
| **60** | NA | 13.85 | NA | 1.66 | 1.85 | 1.85 |
| **120** | NA | NA | NA | **2.07** | **2.45** | **2.28** |
| **Full Exchange** | 18.07 | 18.07 | 18.07 | 5.92 | 5.92 | 5.92 |

**Figure S3.** Comparison of visualization *vs* HX-Express methods for estimation of $t_{1/2}$ of an EX1 process. Both methods are applied to two example peptides (A) 328-351 and (B) 471-478 that exhibit bimodal isotopic distributions during HDX. Visualization method (top): stacked spectra are used to estimate $t_{1/2}$ for both correlated exchange (EX1) and uncorrelated exchange (EX2) in CF4Q, CF4E and CF4Q.A411V arrays. The first spectrum with at least 50% intensity in the high m/z distribution or a symmetric pattern is highlighted in red and labeled #; these represent the visually estimated $t_{1/2}$ of the EX1 process. The first spectrum with at least 50% of full exchange uptake is labeled with green asterisk and represents the visually estimated $t_{1/2}$ of the EX2 process. HX Express method (bottom): HX Express is used to deconvolute the bimodal patterns into two binomial distributions. EX1 (top table): The fractional intensity of the high m/z distribution at each time point is listed; $t_{1/2}$ for EX1 (red) is the time at which this intensity reaches 0.5. EX2 (bottom table): The uptake for the low m/z distribution at each time point is listed; $t_{1/2}$ for EX2 (green) is the time at which this uptake reaches 50% of the full exchange uptake. N.A indicates either missing data or that the two binomial distributions are merged and cannot be deconvoluted. The two methods yield similar estimates; the visualization method has the advantages of being faster and applicable to spectra that lack resolution of the two distributions. Note that the full exchange value was measured in a back exchange control experiment on CF4Q, and assumed to be the same for CF4E and CF4Q.A411V.

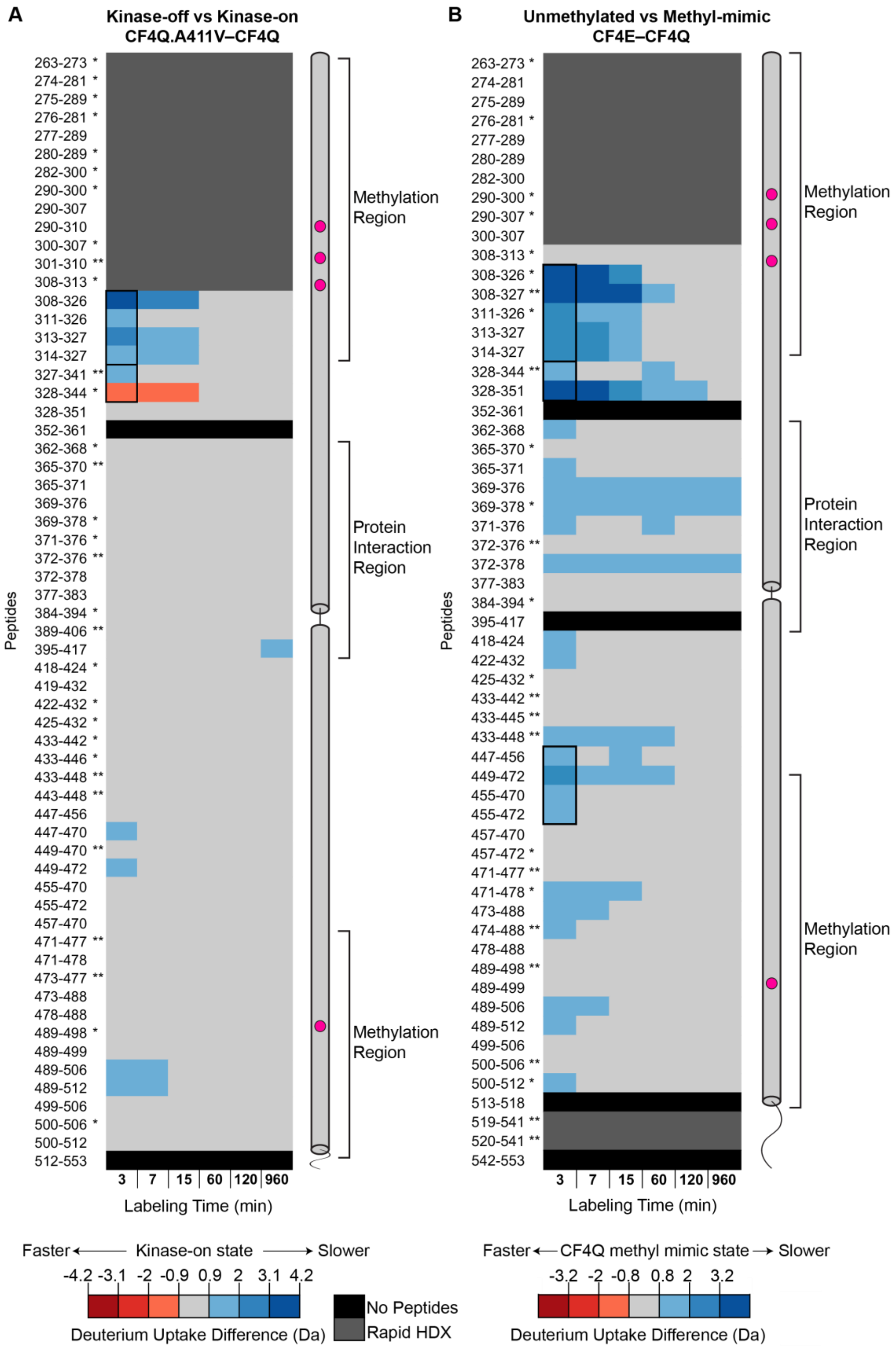
A
Kinase-off vs Kinase-on
CF4Q.A411V–CF4Q
B
Unmethylated vs Methyl-mimic
CF4E–CF4Q
Peptides
Methylation Region
Protein Interaction Region
Labeling Time (min)
3 7 15 60 120 960
Faster ← Kinase-on state → Slower
-4.2 -3.1 -2 -0.9 0.9 2 3.1 4.2
Deuterium Uptake Difference (Da)
No Peptides
Rapid HDX
Faster ← CF4Q methyl mimic state → Slower
-3.2 -2 -0.8 0.8 2 3.2
Deuterium Uptake Difference (Da)

**Figure S4.** Comparison of HDX differences due to signaling state and methylation. For complete set of CF peptides, colors indicate deuterium uptake differences between kinase-on/methyl-mimic (CF4Q) and (A) kinase-off (CF4Q.A411V) or (B) unmethylated (CF4E) states. A subset of the data shown in (A) are presented in Figure 4; a subset of the data shown in (B) are presented in Figure 5. Segments in dark gray undergo very fast exchange (≥ 90% in 3 min), so these HDX experiments do not have the time resolution to detect differences between the two states in these regions. Segments in blue colors undergo slower HDX in the CF4Q kinase-on state; segments in light gray show no significant difference (difference ≤ ±0.9 Da). A single peptide (328-344) displays faster HDX in the CF4Q kinase-on state (red color). We concluded this was not significant beause overlapping peptides 327-341 and 328-351 do not show this change. Single asterisks indicate the peptide is observed in only 3 of the 4 data sets (two replicates for each sample type). Two asterisks indicate the peptide is observed in only 2 of the 4 data sets. In both (A) and (B), black boxes for the 3 min time points enclose overlapping peptides that were averaged to determine the color represented on the structures in Figures 4 and 5.

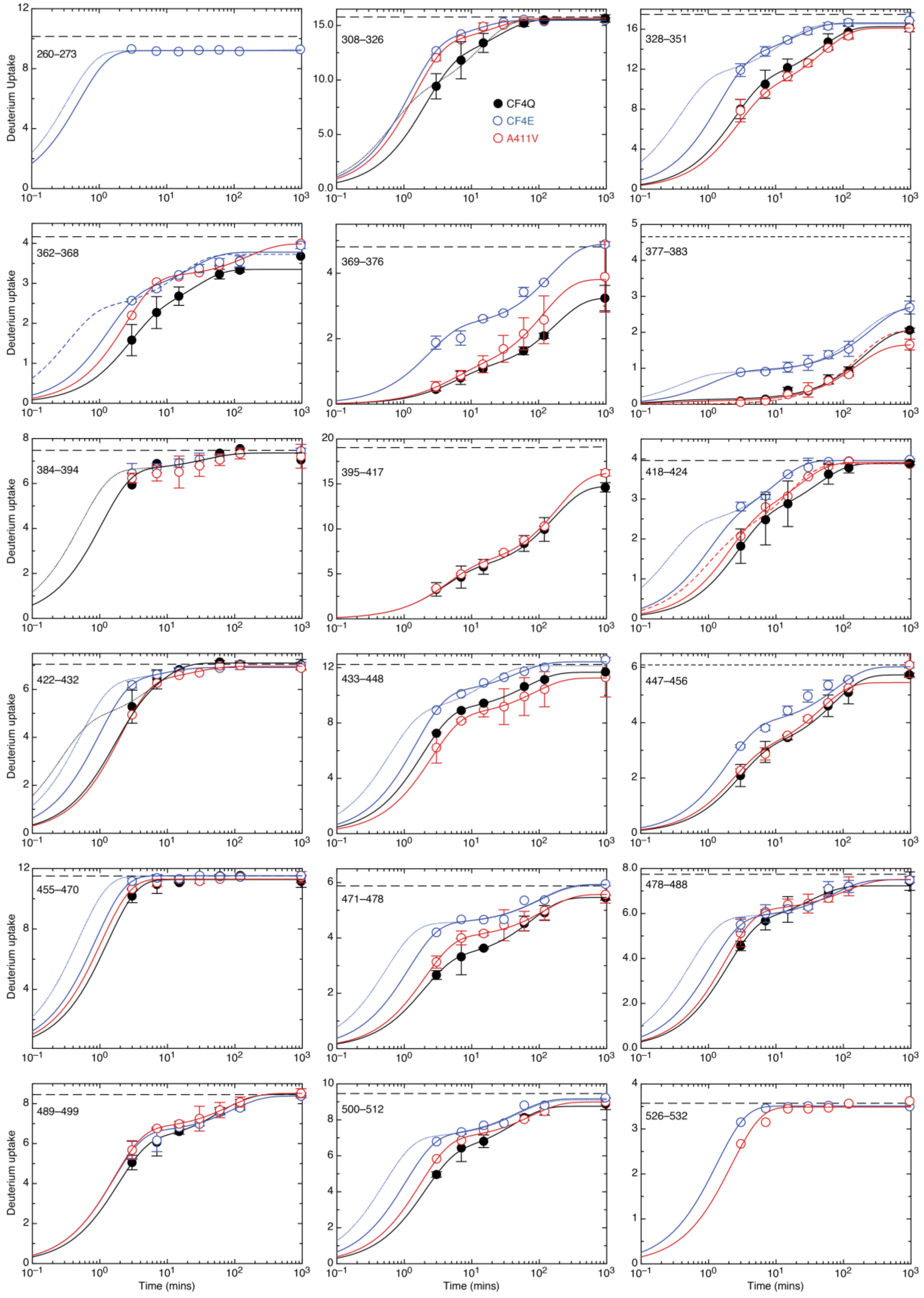

Deuterium Uptake
Deuterium uptake
Time (mins)
CF4Q
CF4E
A411V
260–273
308–326
328–351
362–368
369–376
377–383
384–394
395–417
418–424
422–432
433–448
447–456
455–470
471–478
478–488
489–499
500–512
526–532

**Figure S5.** Uptake plots for representative peptides throughout CF. Curves are best fits of the data to the equation $y = p_1 + p_2 - \left(p_1 e^{-k_1 t} + p_2 e^{-k_2 t}\right)$ for biexponential uptake. Fraction and rate constant of fast ($f_1$, $k_1$) and slow uptake ($f_2$, $k_2$) for each peptide are reported in Table S2. A few peptides were well fit with a monoexponential curve. “Very fast” uptake (already high at 3 min) was fit to the minimum k that would fit the first time point (solid curves) as well as to a faster k (dotted or dashed curve) to demonstrate that the data only determine a lower limit for k (eg $k \geq 0.2\ min^{-1}$), as reported in Table S2. Complete exchange (as measured in a back exchange control) is represented by the horizontal dashed black line near the top of each plot.

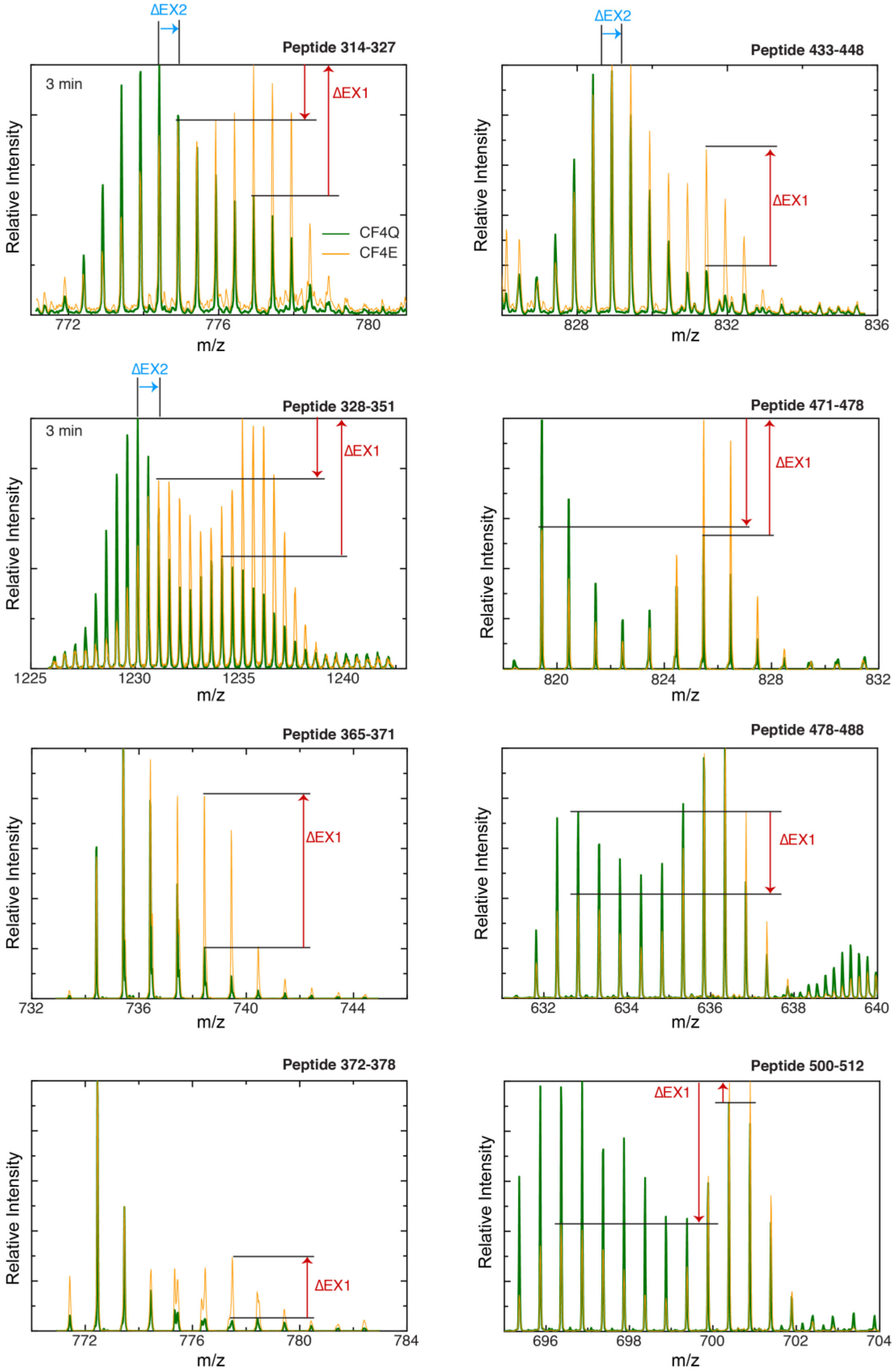
ΔEX2
Peptide 314-327
3 min
ΔEX1
CF4Q
CF4E
Relative Intensity
772
776
780
m/z
ΔEX2
Peptide 433-448
ΔEX1
Relative Intensity
828
832
836
m/z
ΔEX2
Peptide 328-351
3 min
ΔEX1
Relative Intensity
1225
1230
1235
1240
m/z
Peptide 471-478
ΔEX1
Relative Intensity
820
824
828
832
m/z
Peptide 365-371
ΔEX1
Relative Intensity
732
736
740
744
m/z
Peptide 478-488
ΔEX1
Relative Intensity
632
634
636
638
640
m/z
Peptide 372-378
ΔEX1
Relative Intensity
772
776
780
784
m/z
Peptide 500-512
ΔEX1
Relative Intensity
696
698
700
702
704
m/z

**Figure S6.** Superpositions of mass spectra show largest changes are in the fast initial EX1. Mass spectra after 3 min of HDX for CF4Q or CF4E are superimposed for peptides that show a change in HDX and detectable EX1. Spectra are annotated to highlight the change from CF4Q to CF4E in EX2 (blue horizontal arrow indicates shift in the position of the most intense line) and in EX1 (red vertical arrows indicate the decrease in the area of the isotopic distribution at low m/z and the increase in the area of the fully exchanged isotopic distribution). Note that the spectra are arbitrarily scaled to equalize the most intense peak, so the total area of the isotopic distributions for the two samples are not necessarily equal.


1. Wang, X., Vu, A., Lee, K., and Dahlquist, F. W. (2012) CheA-receptor interaction sites in bacterial chemotaxis. *J. Mol. Biol.* **422**, 282–290
2. Piasta, K. N., Ulliman, C. J., Slivka, P. F., Crane, B. R., and Falke, J. J. (2013) Defining a key receptor-CheA kinase contact and elucidating its function in the membrane-bound bacterial chemosensory array: A disulfide mapping and TAM-IDS study. *Biochemistry*. **52**, 3866–3880
3. Briegel, A., Li, X., Bilwes, A. M., Hughes, K. T., Jensen, G. J., and Crane, B. R. (2012) Bacterial chemoreceptor arrays are hexagonally packed trimers of receptor dimers networked by rings of kinase and coupling proteins. *Proc. Natl. Acad. Sci.* **109**, 3766–3771
4. Cassidy, C. K., Himes, B. A., Alvarez, F. J., Ma, J., Zhao, G., Perilla, J. R., Schulten, K., and Zhang, P. (2015) CryoEM and Computer Simulations Reveal a Novel Kinase Conformational Switch in Bacterial Chemotaxis Signaling Department of Physics and Beckman Institute , University of Illinois at Urbana-Champaign , Urbana , IL 61801 , USA Department of Structural Biolo
5. Vu, A., Wang, X., Zhou, H., and Dahlquist, F. W. (2012) The receptor-CheW binding interface in bacterial chemotaxis. *J. Mol. Biol.* **415**, 759–767
6. Pedetta, A., Parkinson, J. S., and Studdert, C. A. (2014) Signalling-dependent interactions between the kinase-coupling protein CheW and chemoreceptors in living cells. *Mol. Microbiol.* **93**, 1144–1155
